# Activation of Pol theta-mediated end joining begins during mitotic commitment

**DOI:** 10.1101/2024.10.03.616590

**Authors:** Connor P. McBrine, Ryan B. Jensen, Megan C. King

**Affiliations:** Department of Cell Biology, Yale School of Medicine, New Haven, CT, 06520, USA; Department of Therapeutic Radiology, Yale School of Medicine, New Haven, CT, 06520, USA; Department of Molecular, Cell and Developmental Biology, Yale University, New Haven, CT, 06511, USA

## Abstract

Polymerase theta-mediated end-joining (TMEJ) is a salvage DNA repair pathway that anneals and ligates microhomologies flanking a DNA double-strand break (DSB) downstream of aborted homologous recombination to ensure DSBs are resolved before chromosome segregation. Here, we use a sensitive deep-sequencing approach to identify DSB repair outcomes at an endogenous locus. We observe that TMEJ is impaired by depletion or inhibition of DNA2, a component of the DSB resection machinery. Mechanistically we find that DNA2 deficiency influences TMEJ efficiency by slowing S-phase progression, thereby preventing cells from reaching mitotic commitment at the G2/M transition, when we find that TMEJ is optimally licensed to occur. Further interrogation of the cell cycle regulation of TMEJ reinforces a strict requirement for the mitotic kinase PLK1 in enabling TMEJ at the G2/M transition. We surprisingly find CDK1 to be largely dispensable for TMEJ, but important for a small subset of repair events likely occurring in mitosis. These findings have important implications for modulation of TMEJ in response to cell cycle kinase inhibitors that are in active development as anti-tumor agents.

## Introduction

DNA double-strand breaks (DSBs) are a threat to genome stability and must be resolved by the cell’s DNA repair machinery. Homologous recombination (HR) is the high-fidelity pathway that fully restores the genetic information of the broken chromosome by synapsing with the intact sister chromatid and using it as a template for new synthesis. However, when HR repair is defective (as is the case with cancers mutated in *BRCA1* or *BRCA2*), a salvage repair pathway called polymerase theta-mediated end-joining (TMEJ) can join the two broken ends of DNA back together to create an intact, but mutated, chromosome [1–4]. During TMEJ, polymerase theta (Polθ, encoded by *POLQ*) scans for and aligns microhomologies between exposed ssDNA tails generated during attempted HR on either side of the DSB [5]. After a flap trimming step, Polθ initiates short-range synthesis that bridges the DSB [6, 7]. TMEJ is inherently error-prone, and the joining of two microhomologies into a single copy will result in the full deletion of sequence between two microhomologies. For this reason, TMEJ is largely synonymous with microhomology mediated end-joining (MMEJ) in mammalian systems [1]. Microhomologies are typically 2-6 nt in length [8]. However, more complex events involving intra-strand snap-back synthesis can also result in insertion products [9, 10]. In either case, evidence of TMEJ usage can be inferred from the sequence characteristics of the insertions/deletions (indels) at a repaired DSB [6, 11].

DSB repair mechanism choice is governed by resection, the 5’ to 3’ nucleolytic degradation at the DSB to generate 3ʹ single-stranded DNA (ssDNA) tails. Non-homologous end-joining (NHEJ) is preferred when resection is inhibited, for example by 53BP1 and the Shieldin complex [12, 13]. In contrast, DSB resection is a shared, initial processing event that lies upstream of both HR and TMEJ [14]. The MRE11-RAD50-NBS1 complex (MRN) together with the pro-HR factor CtIP initiates resection with short-range nucleolytic degradation of the 5’ strands of the DSB. Long-range resection (up to several kilobases) necessary for efficient HR is performed by Exonuclease 1 (EXO1) and/or the Bloom’s syndrome helicase (BLM) acting with the DNA replication helicase/nuclease 2 (DNA2). BLM helicase physically interacts with Topoisomerase IIIα (TOP3α) and the RecQ-mediated genome instability protein 1 (RMI1) and RMI2 to form the BTR complex [15]; the BTR complex stimulates BLM-DNA2-mediated resection *in vitro* but may not be a strict requirement *in vivo* [16–18]. There is little consensus on the division of labor between EXO1 and BLM-DNA2 in promoting the resection necessary for HR, particularly when considering the variability across eukaryotic models. Yeast genetics suggest that the Exo1 nuclease preferentially drives resection while the BLM orthologs *S. cerevisiae* Sgs1 or *S. pombe* Rqh1 acting in concert with Dna2 can compensate in its absence [19–26]. In mammalian systems, some studies also suggest a predominant physiological role for EXO1 [27] while others indicate EXO1 and BLM-DNA2 are fully redundant [28, 29]. Recent work argues that the specific chemistry at a DSB (bulky adducts, ribonucleotides, and damaged bases) may influence the requirement for processing by EXO1 and/or BLM-DNA2, thereby explaining the cell’s need for two (potentially competing) resection pathways [30].

While resection is necessary for the exposure of microhomologies upstream of TMEJ, it is unclear exactly how much resection is required. Although TMEJ can result in large deletions and rearrangements, most characterized repair products involve microhomologies very close (within 15 bp) of the DSB ends [6]. The short-range resection factors MRE11 and CtIP both contribute to TMEJ repair [31–33], while the long-range resection factors EXO1 and BLM have less obvious roles. Indeed, in *S. cerevisiae,* Exo1 and the BLM ortholog Sgs1 are both dispensable for MMEJ repair involving microhomologies immediately flanking the DSB [34]. In human cell lines there are conflicting reports, with depletion of either EXO1 or BLM observed to promote TMEJ in a TMEJ versus HR reporter system [14] while BLM depletion suppresses TMEJ in a TMEJ versus NHEJ reporter system [35]. In *S. cerevisiae,* Dna2 deficiency partially impairs MMEJ [36], although a role for DNA2 in mammalian TMEJ has not been explored.

Further complicating this landscape, both the EXO1 and DNA2 nucleases have additional functions outside of DSB resection. EXO1 has roles in DNA replication through its lagging strand flap removal activity [37] and its resolution of telomeric G-quadruplex DNA [38], but additionally participates in mismatch repair (MMR) and nucleotide excision repair (NER) by generating ssDNA gaps on the damaged strand of DNA [39]. DNA2 has essential roles in Okazaki fragment processing [40] as well as in replication fork restart [41–45], and maintenance of centromeric, telomeric, and even mitochondrial DNA [46, 47]. An essential protein for organism viability, DNA2 is also overexpressed in many cancers [48] and is increasingly being investigated as a drug target whose inhibition can induce replication stress and cell cycle arrest [49–52].

In addition to resection, cell cycle status is thought to play a major role in regulating TMEJ usage. NHEJ is active throughout interphase, while HR and its initial resection step are suppressed during the G1 phase when the sister chromatid has not yet been replicated. TMEJ, acting primarily as a backup for HR, is therefore largely inactive during G1 [14], although there are reports that aberrant resection in G1 leads to the engagement of the TMEJ pathway [53, 54]. More commonly, however, unrepaired G1 DSBs likely persist into S or G2/M phase for eventual repair by HR or TMEJ [55]. Specifically, mitosis has been described as the optimal window when TMEJ is activated as a final failsafe repair pathway before chromosome segregation. The HR factors RAD52 and BRCA2 have been suggested to restrain Polθ activity during S and G2 phases, thereby delaying TMEJ repair until the onset of mitosis [56]. Several factors are proposed to further promote recruitment and activity of Polθ during mitosis, including TOPBP1 [57, 58] and RHINO [59]; both of these mechanisms are further regulated by the Polo-like kinase 1 (PLK1), which directly phosphorylates Polθ [57–59]. Of note, there is a network of kinases including PLK1, Aurora A, and multiple CDK-cyclins (CDK1-CyclinA; CDK2-CyclinA, CDK1-CyclinB) that become activated at the G2/M transition and coordinate mitotic entry and transit through mitosis. These kinases operate in a positive feedback loop to quickly enhance mitotic signaling; in the conventional model, CyclinA-CDK1 first activates Aurora A, which in turn activates PLK1 [60, 61]. Both cyclin-dependent kinase 1 (CDK1) and Aurora Kinase A have been suggested to facilitate TMEJ during mitosis [62].

Inhibition of TMEJ is synthetic lethal with genetic conditions that give rise to HR-deficiency [63]. As such, several Polθ inhibitors are in development to treat HR-deficient cancers, including those that have become resistant to PARP inhibitors [63, 64]. However, further study is needed to understand all of the genetic requirements for TMEJ, and whether factors upstream or downstream of Polθ activation can be alternative drug targets in HR-deficient cancers or biomarkers to predict the Polθ inhibitor response.

Here, we leveraged a sequencing-based approach to interrogate repair products of a Cas9-induced DSB. Initially seeking to investigate how DSB end resection mechanisms modulate the use of TMEJ, our data revealed that shifts in cell cycle progression upon perturbation of resection factors is predominantly responsible for their effect on TMEJ usage. By further dissecting this cell cycle regulation, we discovered that cells activate TMEJ at mitotic commitment, but prior to mitotic entry, suggesting that this salvage repair pathway becomes favored just prior to chromosome condensation and segregation. Thus, while we found that a subset of complex TMEJ repair products take place in mitosis, simple TMEJ repair likely occurs in a burst immediately prior to mitotic entry.

## Results

### Polymerase theta inhibition alters DSB repair outcomes

We first developed a DSB repair assay to study TMEJ in RPE1-hTERT and U2OS cells. RPE1-hTERT is an immortalized but non-transformed cell line that is nearly karyotypically normal and generally regarded as representative of normal cell physiology [65, 66] while U2OS is an osteosarcoma cell line commonly used as a model system for studying DNA repair mechanisms including DSB end resection and HR [27, 28, 67]. We assessed TMEJ proficiency by interrogating the repair outcome at a single, site-specific DSB induced by a Cas9 ribonucleoprotein (RNP) complex delivered by electroporation. The Cas9 RNP is targeted to a region within the *ADGRL2* gene that has previously been reported to be permissive for HR repair of Cas9-induced DSBs [68], suggesting potential for engagement of the TMEJ pathway as well. We evaluated the outcomes 24 hours after Cas9 RNP delivery to allow time for cellular DNA repair mechanisms to resolve the damage. During this 24-hour window we expected many cells to accurately repair the DSB back to its original sequence either through error-free NHEJ or HR, in which case the repaired target sequence is likely to be re-cut by the Cas9 RNP. This cut-repair cycle therefore drives a selective pressure towards mutagenic outcomes [69]. To evaluate the repair events, we purified genomic DNA and PCR amplified the region containing the Cas9 recognition site. Only fully intact (i.e. uncut or repaired) genomes can be amplified; persistent DSBs cannot serve as a PCR template and therefore will not be represented in the dataset. We generated libraries of PCR amplicons spanning the cut site that were sequenced on the Illumina platform to precisely identify repair products. We employed the computational tool SIQ [70] to characterize all sequences according to mutation type (i.e. deletion size, microhomology usage, insertion size, and more complex rearrangements), which then allowed sorting into probable TMEJ or NHEJ events.

Both RPE1 and U2OS cells were able to repair a Cas9-induced DSB in an efficient but mutagenic manner. A representative spectrum of the most common repair products in U2OS (insertions, deletions, deletions with insertions, and deletions with templated insertions) reveals the wide variety of deletion sizes and usage of microhomologies at the repair junctions (Figure 1A). To experimentally define the repair products that rely on Polθ activity we leveraged the Polθ inhibitor (Polθi) ART558. We clearly observed large shifts in the relative abundance of mutation types in cells recovered into ART558 compared to those treated with DMSO after Cas9 transfection, as revealed by the color-coded sequencing outcomes (Figure 1A, Supplemental Figures S1, S2).

**Figure 1.**
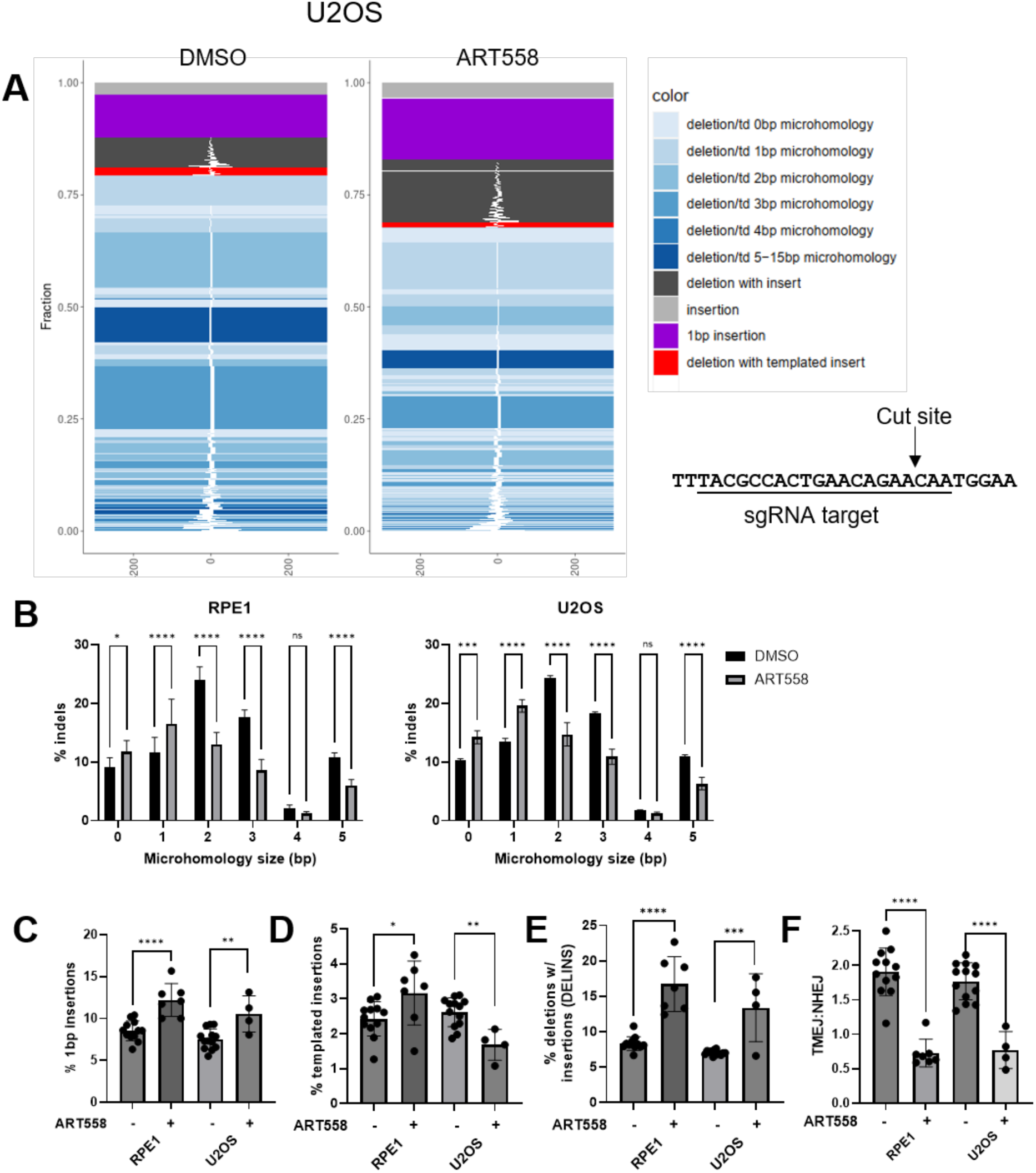
Inhibition of polymerase theta alters DSB repair spectra. **(A)** Representative spectrum of DSB repair events in U2OS cells following cleavage and repair of the depicted Cas9 cut site. Cells were cultured with the Polθi ART558 (10 μM) or DMSO for 24 hours after Cas9 RNP transfection. Genomic DNA was then harvested and sequenced, and mutation types were sorted and visualized with the software tool Sequence Interrogation and Quantification (SIQ). **(B)** ART558 suppresses microhomology usage during end-joining repair. Deletion events in RPE1 and U2OS cells treated with DMSO vs ART558 (10 μM) were sorted by the size of microhomology on either side of the cut site utilized to join the DSB. The frequency of microhomologies above 5 bp was negligible. Significance was determined by one-way ANOVA with Šídák’s multiple comparison test. **(C-E)** ART558 treatment alters distinct mutation events. Graphs depict **(C)** % 1 bp insertions, **(D)** % templated insertions (TINS), and **(E)** % deletions with insertions (DELINS) as a percent of total indels in RPE1 and U2OS cells transfected with Cas9 RNP and recovered into ART558 (10 μM). 1 bp insertions are a hallmark of NHEJ and TINS are a hallmark of TMEJ. Significance for each pair was determined by unpaired, two-tailed t-test. **(F)** A TMEJ:NHEJ summary metric was calculated as the ratio of (≥ 2 bp deletions) / (≤ 1 bp deletions + 1 bp insertions). **(C-F):** Datapoints depict independent biological experiments with mean ± SD. * p ≤ 0.05, ** p ≤ 0.01, *** p ≤ 0.001, **** p ≤ 0.0001.

Looking into more detail, we observed that ART558 treatment increased end-joining deletions utilizing 0 or 1 bp of homology, but suppressed usage of microhomologies ≥ 2 bp, in both RPE1 and U2OS cells (Figure 1B). ART558 treatment also caused characteristic shifts in rare, but highly informative mutation events. Insertions of 1 bp (typically hallmarks of canonical NHEJ) were elevated by ART558 treatment in both cell lines (Figure 1C). Conversely, templated insertions (“TINS”, associated with TMEJ usage [71, 72]) were only suppressed by ART558 treatment in U2OS cells. In RPE1 cells TINS were unexpectedly elevated (Figure 1D), although the reason for this difference is not immediately clear. Lastly, we observed that ART558 treatment significantly increased deletions with insertions (“DELINS”) that do not involve microhomology usage (Figure 1E). To capture this information in a single metric, we adopted a ratiometric measurement that facilitates comparisons of the relative TMEJ utilization across experiments. Since the templated insertions effects were inconsistent across cell lines, and DELINS lack a clear mechanistic basis for TMEJ, we chose to define the ratio of TMEJ to NHEJ as (≥ 2 bp deletions) / (≤ 1 bp deletions + 1 bp insertions) (Figure 1F).

### Depletion of DNA2 but not EXO1 impairs TMEJ repair

Having developed a deep mutational analysis assay, we next tested how siRNA knockdown of the canonical long-range resection factors EXO1 and DNA2 affects TMEJ usage. We confirmed near-complete knockdown of the target transcript levels by 48 hours (Supplemental Figure S3A, B). We observed that depletion of DNA2 (but not EXO1) caused a marked reduction in TMEJ repair events compared to NHEJ events (Figure 2A). To take an orthogonal approach we leveraged the DNA2 inhibitor (DNA2i) C5, which selectively binds to an allosteric pocket in the helicase domain of DNA2, suppressing its nuclease, ATPase, and helicase activities. We observed similar TMEJ defects when treating the cells with the DNA2i for 24 hours following Cas9 RNP transfection (Figure 2B, Supplemental Figures S1, S2). DNA2i treatment increased 1bp insertions and suppressed templated insertions, indicative of a defect in TMEJ activity (Supplemental Figure S3C) although there are subtle differences in the pattern between the DNA2i and siDNA2 treatment conditions (Supplemental Figure S3C-E, S4).

**Figure 2.**
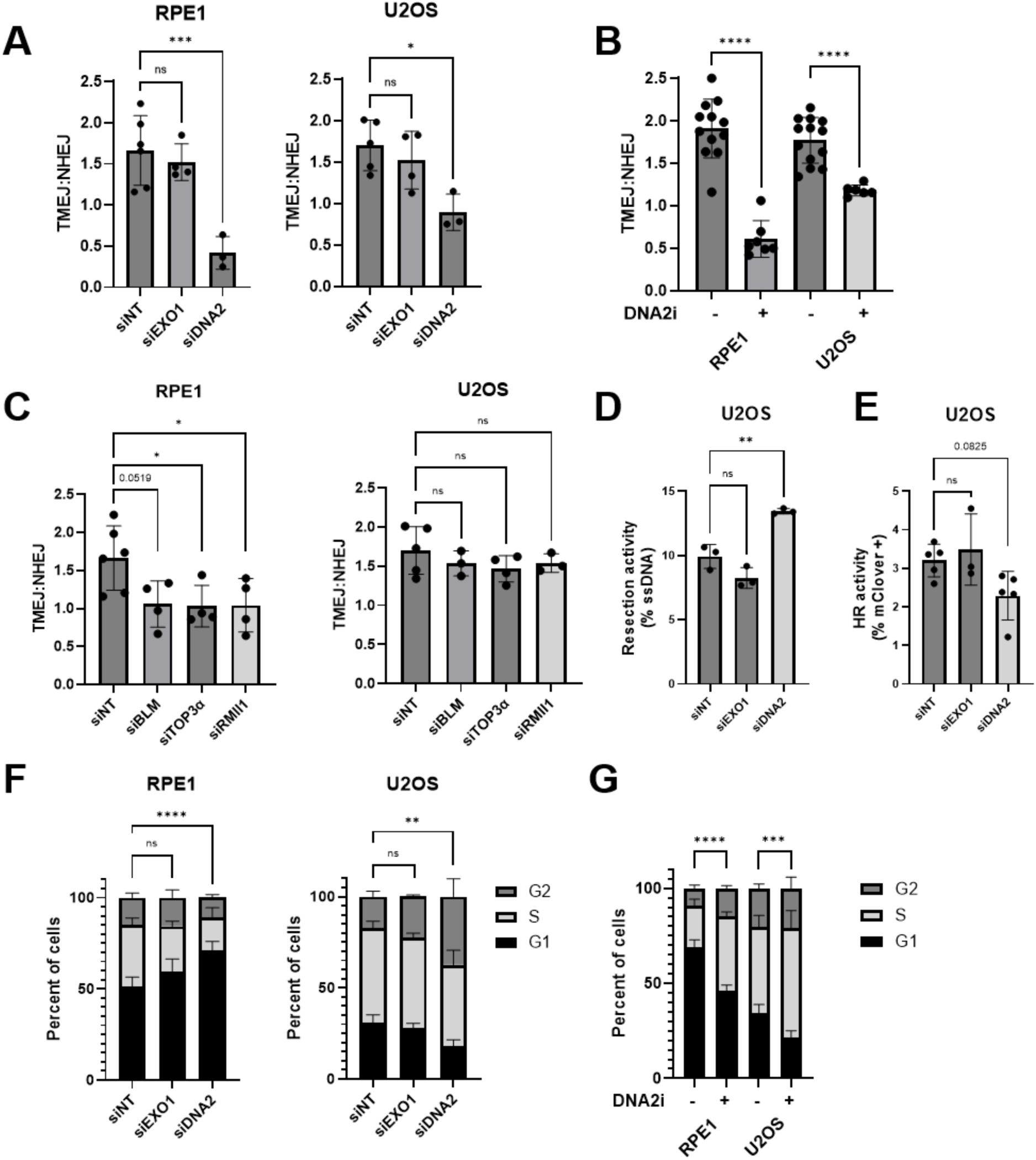
DNA2 depletion impairs TMEJ, alters DSB resection, and perturbs the cell cycle. **(A)** Knockdown of DNA2 suppresses TMEJ. TMEJ vs NHEJ repair events of a Cas9-induced DSB in RPE1 and U2OS cells depleted of EXO1 or DNA2 for 48 hours prior to Cas9 RNP transfection. Significance was determined by one-way ANOVA with Šídák’s multiple comparison test. **(B)** Inhibition of DNA2 by recovery into the DNA2 inhibitor C5 (60 μM) for 24 hours after transfection suppresses TMEJ. Significance for each pair was determined by unpaired, two-tailed t-test. **(C)** The BTR complex appears dispensable for TMEJ. TMEJ vs NHEJ repair events of a Cas9-induced DSB in RPE1 and U2OS cells depleted of the BTR complex proteins BLM, TOP3α, and RMI1. Significance was determined by one-way ANOVA with Dunnett’s multiple comparison test. **(D)** DNA2 depletion enhances resection. Resection 8 hours post-DSB induction at a site 1040 bp distal to the Cas9-induced DSB in U2OS cells following 48 hours of siRNA knockdown of the indicated gene products. Significance was determined by one-way ANOVA with Šídák’s multiple comparison test. **(E)** Neither EXO1 nor DNA2 depletion significantly impairs HR efficiency of LMNA-mClover repair assay in U2OS cells treated with the indicated siRNAs. Significance was determined by one-way ANOVA with Dunnett’s multiple comparison test. **(F)** DNA2 knockdown causes G1 accumulation in RPE1 cells and S/G2 accumulation in U2OS cells. Cell cycle was determined by propidium iodide staining and flow cytometry of RPE1 and U2OS cells treated for 48 hr with the indicated siRNAs. Significance was determined by one-way ANOVA with Šídák’s multiple comparison test on the %G1 value for each sample. **(G)** Pharmacological inhibition of DNA2 causes S/G2 accumulation. Cell cycle was determined by propidium iodide staining and flow cytometry of RPE1 and U2OS cells treated for 24 hr with DNA2i (60 μM C5). Significance was determined by unpaired, two-tailed t-test on the %G1 value for each sample. **(A-G):** Data are the mean ± SD of at least 3 independent biological replicates. * p ≤ 0.05, ** p ≤ 0.01, *** p ≤ 0.001, **** p ≤ 0.0001, ns = not significant.

We initially hypothesized that the decrease in TMEJ usage in the absence of DNA2 activity could reflect DNA2’s role in DSB resection upstream of TMEJ, preventing TMEJ as a possible downstream repair option. DNA2 is reported to cooperate with BLM helicase during DSB resection, potentially involving the full BTR complex of BLM-Topoisomerase IIIα-RMI1-RMI2 [16–18]. We therefore investigated if siRNA-depletion of the BTR complex (Supplemental Figure S3A, B) phenocopies loss of DNA2 function.

In RPE1 cells, siRNA depletion of TOP3α and RMI1 caused a modest but statistically significant reduction in TMEJ usage, with the TMEJ impairment upon BLM depletion trending in a similar fashion (Figure 2C). In contrast, none of the individual siRNA depletions altered the TMEJ:NHEJ ratio in U2OS cells (Figure 2C). Although hallmark 1bp insertions were also slightly elevated in RPE1 cells treated with siBLM and siTOP3α (suggesting a shift away from TMEJ to NHEJ), all other 1 bp insertion and templated insertion frequencies were unaffected (Supplemental Figure S3D, E, S4B). Taken together, our observations suggest that TMEJ is slightly sensitive to BTR complex inhibition in RPE1 cells but not U2OS cells despite the requirement for DNA2 activity in both cell types (Figure 2 A-B). Given this lack of concordance, we began to suspect that BTR-dependent, DNA2-mediated resection may not explain why DNA2 inhibition had such a major effect on TMEJ.

### Resection and HR status do not account for DNA2’s effect on TMEJ

To begin to address whether the effects of disrupting DNA2 activity on TMEJ reflects a potential contribution to its canonical roles in DSB processing, we next interrogated DSB end resection directly. Although a number of indirect assays to measure DSB resection have been used broadly (e.g. the recruitment of RPA to ssDNA including that generated by resection, giving rise to nuclear foci), we sought to more explicitly measure the resection status of the specific Cas9-induced DSB that we evaluated for TMEJ repair. To that end, we employed a qPCR-based resection assay [67]. We extracted genomic DNA at various points after DSB induction and subjected it to *in vitro* ApoI digestion, which is a frequent cutter of dsDNA but does not cleave ssDNA (i.e. resected DNA), which we subsequently quantified by qPCR (Supplemental Figure S5A). We probed a region 1040 bp distal to the DSB, which is well beyond the range that MRN-CtIP can resect [73–75], and therefore requires the long-range machinery of EXO1 and/or BLM-DNA2 for resection. We found that the percent of DSBs that are resected gradually increased to a maximum at 12 hours post-transfection, returning to baseline at 24 hours, suggesting that most DSBs had been repaired (Supplemental Figure S5B). At 24 hours post-transfection we found over 65% of DSBs had been repaired with mutagenic indels, indicating high transfection efficiency and robust DSB induction (Supplemental Figure S5C).

Applying this resection assay to U2OS cells, we observed that EXO1 depletion had little effect, whereas DNA2 depletion surprisingly increased DSB resection activity (Figure 2D). This would imply firstly that DNA2 is not strictly required for DSB resection, or at least not required for long-range resection on the order of ∼1kb at this specific euchromatic locus. Secondly, we conclude that DNA2’s effect on TMEJ is unlikely to be related to its direct roles in DSB resection. An explanation for the increase in resection in siDNA2 cells over the siNT control baseline is not immediately obvious. While it is plausible that EXO1-dependent resection could be up-regulated upon DNA2 depletion, we suspect instead that a shift in cell cycle distribution towards S/G2 phase when resection is licensed may be the root cause (see more below).

We next evaluated the canonical repair pathway downstream of DSB resection, HR. Since TMEJ may act as a back-up for failed HR repair, we considered that elevated HR function could also diminish apparent TMEJ usage, and vice versa. It was not possible, however, to infer HR function from the sequencing results of our model DSB system because perfect HR repair is indistinguishable from error-free NHEJ or uncut genomes. We therefore turned to the established LMNA-Clover homology-directed repair (HDR) assay that measures HR repair of a Cas9-induced DSB within the *LMNA* gene. The induced DSB is repaired using an exogenous plasmid donor containing the mClover reporter gene (plus upstream and downstream homology arms) [76]. We observed that neither EXO1 nor DNA2 knockdown significantly reduced the fraction of mClover positive cells (Figure 2E), although there was a subtle down-trend in cells treated with siDNA2. Thus, changes to HR efficiency are unlikely to drive effects on TMEJ usage in this context. We therefore sought other explanations for the TMEJ defects in DNA2-deficient cells.

### DNA2 deficiency alters the cell cycle

A major established contributor to DSB repair mechanism choice is the cell cycle, which exclusively favors NHEJ in G1 but is permissive to HR in S/G2 [77–79]. Recent studies further indicate that TMEJ, considered to be a mutagenic salvage repair pathway, requires transit into mitosis [56, 57, 59, 62], suggesting that it is employed only after the other mechanisms have failed and the G2/M checkpoint has been overcome. Supporting changes in cell cycle progression upon loss of DNA2 activity, we observed modest defects in cellular proliferation upon disruption of DNA2 via siDNA2 or DNA2i treatments (Supplemental Figure S6A, B). Using propidium iodide (PI) staining to measure DNA content by flow cytometry, we next assessed the effect of siDNA2 treatment on the cell cycle. In both the RPE1 and U2OS cell lines, EXO1 depletion had no significant effect on cell cycle distribution (Figure 2F). In contrast, in RPE1 cells we noticed a striking accumulation of siDNA2 cells in the G1 phase of the cell cycle at the expense of the S and G2 populations (Figure 2F). In U2OS cells, DNA2 knockdown caused an increased G2 population at the expense of the G1 population, in line with previous reports [41, 80, 81] (Figure 2F). Pharmacologically, we found that incubation with DNA2i caused an accumulation of S/G2 phase cells at the expense of G1 phase in both RPE1 and U2OS cells (Figure 2G). Taken together, our observations are consistent with the important contribution of DNA2 to efficient DNA replication, as described previously [50, 81].

### DNA2 deficiency impairs S-phase progression and may account for altered DNA repair pathway choice

To further define precisely when DNA2 function is required for normal cell cycle progression we synchronized cells at the G1/S transition with serum-starvation followed by a release into complete media with DNA2i for 24 hours (Figure 3A). We performed this DNA2i treatment in combination with the PLK1 inhibitor BI-2536, thereby forcing cells that successfully completed S-phase to arrest at the G2/M boundary (rather than progressing into the next cell cycle and potentially being mistaken for unreplicated cells in our analysis). Release into DNA2i did not substantially affect viability (Supplemental Figure S6B), but slowed S-phase progression with many cells failing to reach G2 within 24 hours (Figure 3B-E). Indeed, we observed that 64% of mock-treated RPE1 cells reached the PLK1i-induced G2/M arrest within 24 hours of release versus less than 3% of cells also treated with DNA2i (Figure 3B). In U2OS, 74% of PLK1i-treated cells reached G2/M arrest versus just 39% of cells also treated with DNA2i (Figure 3D). The population of slow to replicate cells treated with the DNA2i is also appreciated in the PI histograms (Figure 3C, E). Consistent with the more severe DNA2i-sensitivity in RPE1 cells, most of the replicating cells were still in early S-phase based on the PI histogram, whereas many U2OS cells had reached late S-phase (Figure 3C, E).

**Figure 3.**
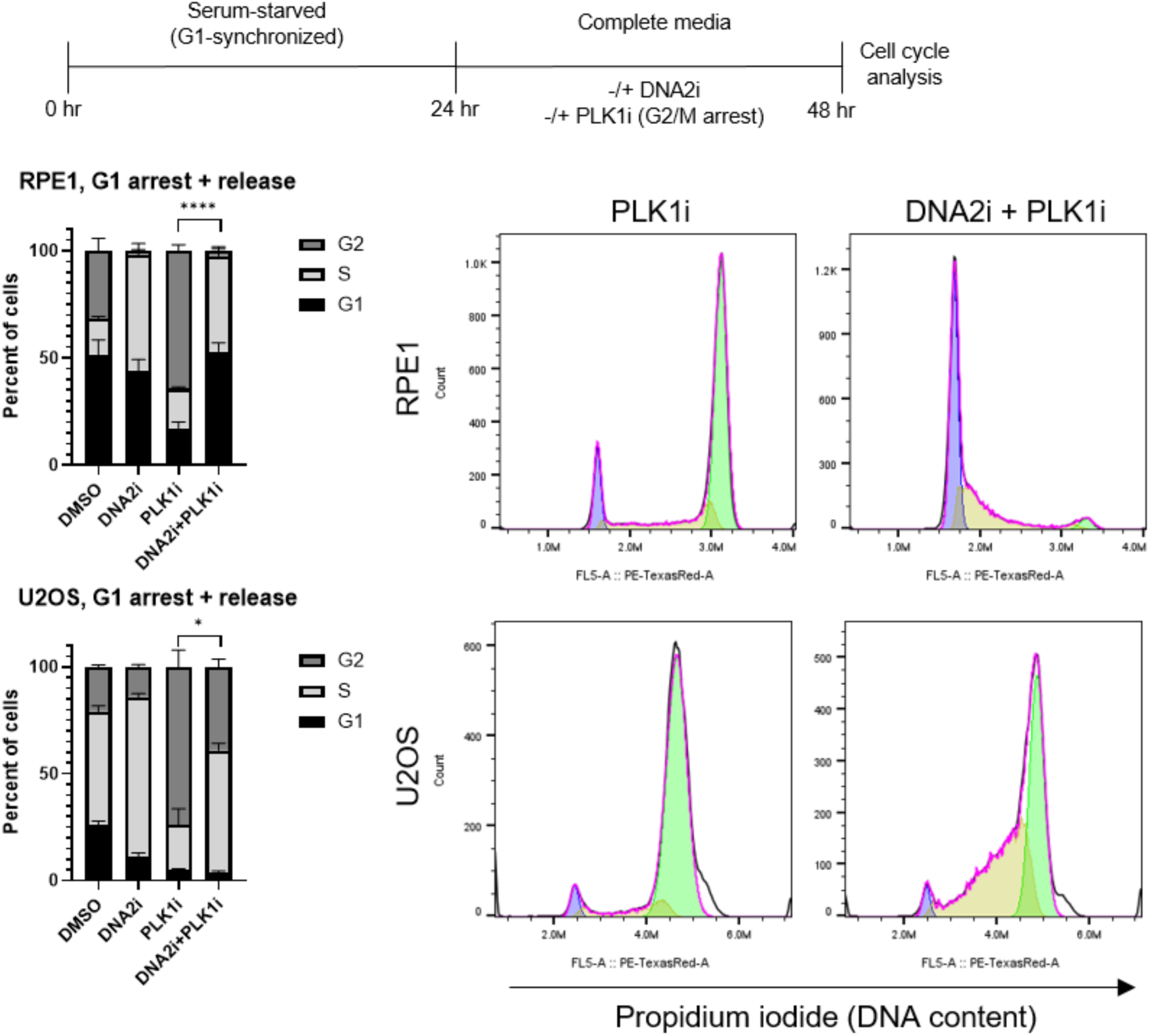
DNA2 inhibition antagonizes cells from reaching the G2/M transition. **(A)** Experimental design of cell cycle arrest and drug treatments. **(B-E)** Inhibition of DNA2 leads to S-phase arrest. Cell cycle analysis of RPE1 **(B,C)** and U2OS **(D, E)** cells G1-arrested with 24 hr serum-starvation and released into complete media containing the indicated drugs for 24 hours. **(B,D)** Data are mean ± SD, n = 3 independent biological replicates. Significance was determined by unpaired, two-tailed t-test on %G2 populations. **(C, E)** Representative histograms of propidium iodide signal show G0/G1 cells in blue, S-phase in yellow, and G2/M cells in green. DNA2i = 60 μM C5, PLK1i = 0.1 μM BI-2536. * p ≤ 0.05, **** p ≤ 0.0001.

We next investigated whether DNA2-deficient cells were simply completing DNA replication slower, or if cell cycle checkpoints that sense DNA damage and replication stress could be mediating an S-phase arrest. To do this, we utilized RPE1 cells with and without the checkpoint master regulator, p53. We observed that RPE1 p53^-/-^ cells are not sensitive to DNA2 knockdown and have the same cell cycle distributions as their untreated counterparts (Supplemental Figure S6C). In contrast to the siDNA2 effects on asynchronous cells, synchronized RPE1 cells released into S-phase with DNA2i failed to achieve productive replication, regardless of p53 status (Supplemental Figure S6D). We suspect this difference arises from the slower loss of DNA2 function in an siRNA knockdown that perhaps permits some replication, whereas acute DNA2 inhibition as cells enter S-phase is a potent inducer of replication stress independent of p53 status.

We considered ATR as a plausible checkpoint protein that could be mediating arrest in DNA2i-treated cells. The ATR kinase is a DNA damage sensor that recognizes RPA-coated ssDNA tracts and arrests replication via CHK1 signaling [82]. However, in synchronized cells released into drug treatments, ATR inhibition failed to rescue the potent replication defects induced by DNA2i (Supplemental Figure S6D, E). As DNA2i-mediated arrest does not seem to act through ATR, we suspect that the replication stress induced by DNA2i treatment is independent of long replication gaps of RPA-coated ssDNA [83]. Instead, DNA2’s role in lagging strand synthesis may be impaired, leading to an accumulation of small, 5’ flaps arising from unprocessed Okazaki fragments [40, 84]. Consistent with this hypothesis, we found that deletion of 53BP1 partially rescues DNA2i-mediated arrest in U2OS cells (Supplemental Figure S6F). Recent work implicates 53BP1 in Okazaki fragment processing [85], and deletion of its ortholog Rad9 in *S. cerevisiae* suppresses the inviable phenotype of *dna2* deletion [86], suggesting that this effect could reflect a conserved function of DNA2.

### TMEJ is licensed by PLK1 activation, likely at mitotic commitment

We next sought to address the apparent paradox in U2OS cells wherein DNA2-deficiency increases resection efficacy at the population level yet decreases TMEJ usage (Figure 2A, B, D). By one line of reasoning, slowed replication will result in more S/G2 cells that preferentially perform resection and thereby generate more of the initial substrate for TMEJ (i.e. resected DSBs). However, contrary to this interpretation, our siDNA2 observations suggest that resection proficiency and S/G2 status alone does not enable TMEJ, since this treatment window (48 hr of siDNA2) results in 82% of cells in S/G2 (vs 69% in untreated cells), yet we observe a 47% reduction in TMEJ compared to untreated cells (Figure 2A, F). Likewise, DNA2i-treated U2OS cells are enriched in the S/G2 phases of the cell cycle yet display poor TMEJ efficiency (Figure 2B, G).

To help address this paradox, we again used serum-starvation and release into drug treatments as a tool, but this time to interrogate DSB repair outcomes. After 24 hours of either asynchronous culture or serum-starvation to induce G1 synchronization, the Cas9 RNP was transfected and all cells were recovered into complete media to allow cells to proceed through the cell cycle in the presence of various drug treatments. We expected that cells released from serum-starvation would exhibit elevated TMEJ usage since they first encounter the Cas9 RNP during S-phase which, according to the accepted model in the field, is when resection can occur. Surprisingly, we instead observed that release from serum-starvation resulted in significantly reduced TMEJ usage (Figure 4A, compare DMSO conditions). As before, DNA2i treatment significantly impaired TMEJ, both in asynchronous and synchronized cells (Figure 4A).

**Figure 4.**
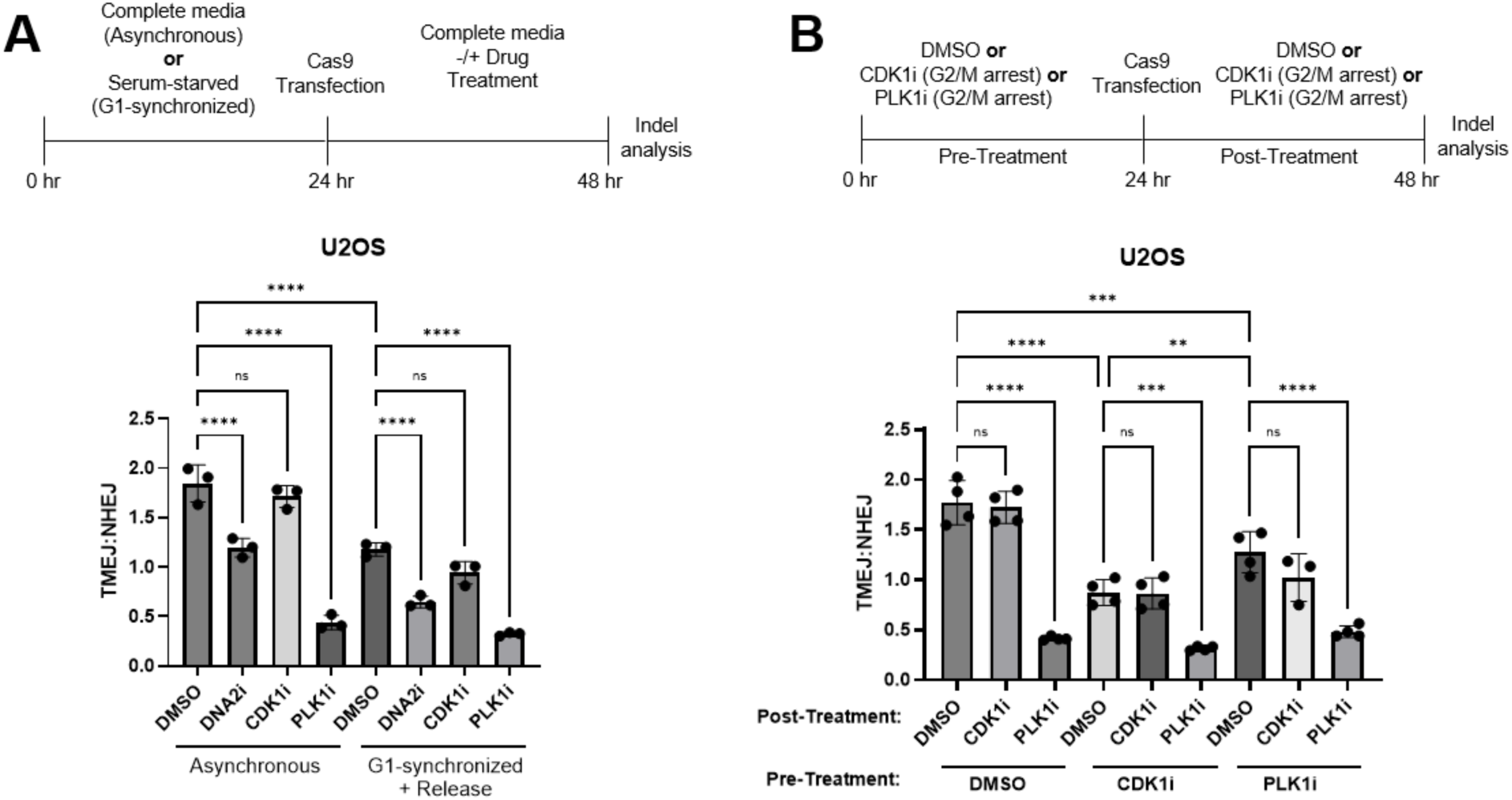
Efficient TMEJ repair requires PLK1 activity, but not CDK1. **(A)** PLK1 activity is required to license TMEJ, while CDK1 is dispensable. TMEJ vs NHEJ repair events in U2OS cells either cultured asynchronously in complete media or synchronized at the G1/S with serum-starvation for 24 hours before Cas9 RNP transfection. After transfection, cells were recovered into complete media containing the indicated drug treatments for 24 hours and then harvested. Data are mean ± SD, n = 3 independent biological replicates. Significance was determined by one-way ANOVA with Šídák’s multiple comparison test. **(B)** TMEJ vs NHEJ repair events of U2OS cells either cultured asynchronously for 24 hours or arrested at the G2/M transition with CDK1i or PLK1i treatment prior to Cas9 RNP transfection (Pre-Treatment). After transfection, cells were recovered into DMSO, CDK1i, or PLK1i treatments (Post-Treatment). Data are mean ± SD, n = 4 independent biological replicates. Significance was determined by one-way ANOVA with Šídák’s multiple comparison test **(A-B)**: DNA2i = 60 μM C5, CDK1i = 5 μM RO-3306, PLK1i = 0.1 μM BI-2536. ** p ≤ 0.01, *** p ≤ 0.001, **** p ≤ 0.0001, ns = not significant.

To provide further insight, we next took advantage of pharmacological inhibitors of mitotic entry. Interestingly, although inhibition of either the CDK1 or the PLK1 kinases is expected to arrest cells at the G2/M transition, we observed significant differences in TMEJ usage in cells treated with the CDK1 inhibitor RO-3306 versus the PLK1 inhibitor BI-2536. While CDK1i had no effect on overall TMEJ usage, PLK1i severely impaired TMEJ (Figure 4A, Supplemental Figures S2, S4A). This effect appears to be generalizable as recovery into PLK1i (but not CDK1i) also impairs TMEJ in RPE1 cells (Supplemental Figures S1, S4A, S7A). While the finding that PLK1 activity is required for a crucial subset of TMEJ repair is expected given several prior studies linking its activation to licensing of TMEJ in mitosis [56, 57, 59, 62], the observation that CDK1 is dispensable is surprising, particularly as CDK1 often primes substrates for phosphorylation by PLK1 [87, 88] and therefore might be expected to act upstream of PLK1. Moreover, as CDK1i prevents mitotic entry, it was puzzling how TMEJ, suggested to take place in mitosis, was unaffected by this treatment.

To further probe these startingly different effects, we transfected the Cas9 RNP into U2OS cells either pre-arrested at the G2/M transition with CDK1 or PLK1 inhibition (pre-treatment) and subsequently recovered them into complete media containing DMSO, CDK1i, or PLK1i (post-treatment). As before, asynchronous cells (DMSO pre-treatment) recovered into PLK1i had impaired TMEJ, but cells recovered into CDK1i did not (Figure 4B). However, we observed that a CDK1i arrest followed by release into DMSO at the time of RNP transfection severely impaired TMEJ (Figure 4B, reduced 51% compared to untreated cells). This suggests that cells first encountering the RNP during mitosis (and then proceeding into the following G1 phase) are not especially permissive for TMEJ; the residual TMEJ activity after release is again CDK1-independent (Figure 4B). By contrast, while PLK1i addition upon inducing the DSB strongly inhibited TMEJ across conditions (Figure 4A, B), release from PLK1i arrest at the time of Cas9 RNP transfection and recovery into DMSO only modestly impaired TMEJ (Figure 4B, pre-treatment with PLK1i, DMSO condition; TMEJ reduced 28% compared to DMSO pre-treatment). Thus, the PLK1i arrest appears to precede the optimal window of TMEJ usage, leading to enhanced TMEJ in the PLK1i pre-treatment compared to the CDK1i-pretreatment condition. In addition, release from PLK1i into CDK1i is permissive to TMEJ (Figure 4B, PLK1i-pre-treatment; CDK1i condition), suggesting that TMEJ can take place after cells recover from the PLK1i-induced arrest even if they do not enter mitosis. Taken together, we conclude that 1) PLK1 activation lies upstream of TMEJ, in line with prior studies [57, 59]; 2) CDK1 is largely dispensable for TMEJ; and 3) PLK1 may license TMEJ in late G2 when cells have committed to, but have not yet entered, mitosis.

### TMEJ depends on Aurora A activity but not mitotic spindle assembly

To further interrogate a model in which cells released from a CDK1i arrest into mitosis have already progressed through the permissive window in which TMEJ is licensed, we next sought to take advantage of an alternate approach to arrest cells in mitosis. As for CDK1 inhibition, when we recovered transfected cells into nocodazole to block mitotic spindle assembly we observed no effect on the TMEJ:NHEJ ratio (Figure 5A). However, also as for CDK1i pre-treatment, Cas9 transfection concurrent with releasing them from a nocodazole block severely impaired TMEJ (Supplemental Figure S7B). Nocodazole treatment therefore behaves exactly like CDK1i treatment in both scenarios, indicating that TMEJ predominantly occurs prior to full CDK1 activation and mitotic spindle assembly in mitosis.

**Figure 5.**
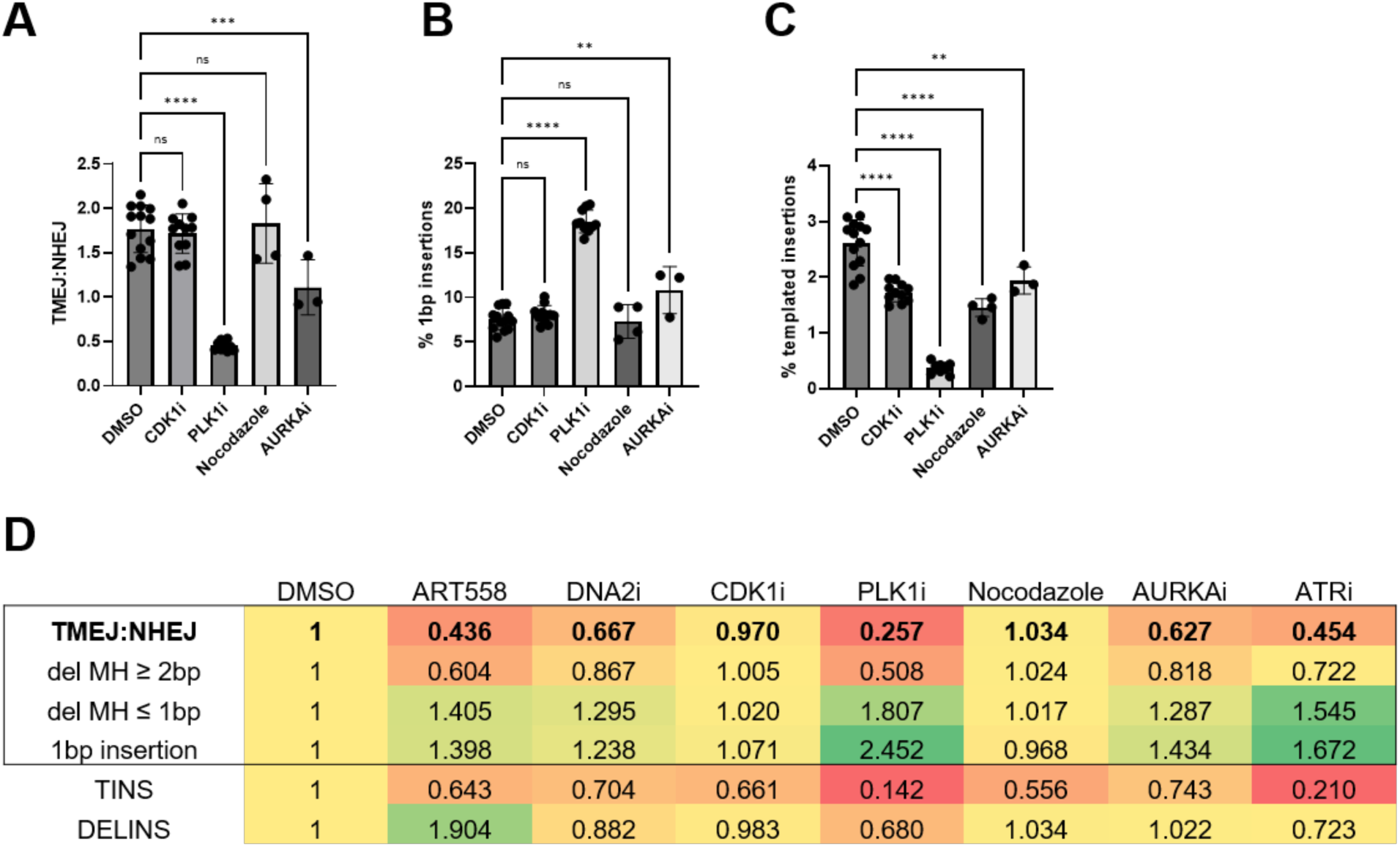
CDK1i and Nocodazole treatments suppress templated insertion events. **(A-C)** DSB repair events of U2OS cells cultured asynchronously and recovered into the indicated drug treatments after Cas9 RNP transfection. Graphs depict the **(A)** TMEJ:NHEJ ratio, **(B)** % 1 bp insertions, and **(C)** % templated insertions. Data are mean ± SD of at least 3 independent biological replicates. Significance was determined by one-way ANOVA with Šídák’s multiple comparison test. **(D)** Summary heatmap of DSB repair events in U2OS cells incubated with the indicated drug treatments for 24 hours after Cas9 RNP transfection. For simpler comparison, all quantities are normalized to the DMSO average for the given mutation type/metric. The normalized TMEJ:NHEJ ratio and the three events that comprise it are boxed; TINS (templated insertions) and DELINS (deletions with insertions) are not incorporated into this ratio but are nevertheless highly informative. **(A-D)** 10 μM ART558, DNA2i = 60 μM C5, CDK1i = 5 μM RO-3306, PLK1i = 0.1 μM BI-2536, 0.2 μM Nocodazole, AURKAi = 1 μM Alisertib, ATRi = 0.5 μM VE-822 in U2OS. ** p ≤ 0.01, *** p ≤ 0.001, **** p ≤ 0.0001, ns = not significant.

Our observations suggest that the optimal window for most TMEJ may be quite small, sometime between the onset of PLK1 and full CDK1 activation. If this is the case, our observations are at odds with the canonical model in which PLK1 and CDK1 activation occur together in a positive feedback loop at mitotic commitment, with CyclinA-CDK1 first acting upstream to enable PLK1 phosphorylation by Aurora Kinase A [61, 89]. However, a more nuanced model describes PLK1 becoming gradually activated towards late G2 phase in a signaling cascade dependent upon CyclinA-CDK1/2, Aurora Kinase A, and Bora. PLK1 subsequently acts via Cdc25C1 to activate CyclinB1-CDK1 and commit to mitotic entry [90–93]. Thus, we considered that an early wave of PLK1 activity might license TMEJ by phosphorylating Polθ (and possibly its cofactor RHINO)[57, 59] during mitotic commitment independent of CDK1. In such a model, the Aurora A kinase would be expected to play an important role in licensing TMEJ. Indeed, we observed a defect in TMEJ, albeit more modest than that maximally observed in the presence of PLK1i, when recovering transfected cells into the Aurora Kinase A (AURKA) inhibitor Alisertib (Figure 5A). Taken together, these observations suggest that the optimal window for TMEJ occurs at mitotic commitment in late G2 rather than in mitosis.

### Templated insertions require transit into mitosis

To gain additional insight, we again looked in greater detail for telltale mutation events indicative of the TMEJ and NHEJ balance. As expected, the diminished TMEJ:NHEJ ratio when recovering into PLK1i or AURKAi was corroborated by an increase in 1 bp insertions and a decrease in templated insertions (Figure 5A-C, Supplemental Figure S2). As neither CDK1i nor nocodazole treatment altered the TMEJ:NHEJ ratio, 1 bp insertions were accordingly also unchanged. Surprisingly, however, the frequency of templated insertions was significantly decreased across all inhibitor treatments (Figure 5C). Templated insertions are complex mutation events that arise when microhomologies anneal either on the opposite side of the DSB (in *trans,* leading to direct repeats) or on the same strand after intramolecular hairpin formation (in *cis*, leading to inverted repeats). Polθ primes synthesis from these annealed substrates before eventually aborting synthesis and restarting a second round of microhomology annealing and synthesis [6, 71]. A partial schematic of the formation of a templated insertion event common in our datasets is provided as an example (Supplemental Figure S8).

To directly compare mutation type frequency across all inhibitor treatments used throughout our study, we normalized all metrics to the DMSO control (Figure 5D, Supplemental Figure S9). In U2OS cells, the TMEJ:NHEJ ratio and all mutation events comprising TMEJ or NHEJ are not affected by recovery into CDK1i or nocodazole. However, templated insertions are substantially reduced. These results suggest that a small subset of mutation events attributable to TMEJ may indeed require CDK1 activity and transit into mitosis itself, perhaps even occurring downstream of spindle assembly.

Integrating these insights, we can now provide a model for why impaired TMEJ upon DNA2 deficiency is best explained by its slow replication phenotype. Whether by siRNA depletion in asynchronous cells, or release into S-phase with DNA2i, DNA2-deficiency gives rise to fewer cells that reach the critical window of TMEJ activity near the G2/M transition. We can thereby reconcile the apparent paradox that DNA2 deficiency results in a larger fraction of S/G2 cells and elevated resection proficiency in U2OS cells, yet fails to efficiently engage the TMEJ repair pathway – the cells simply fail to reach the G2/M transition. More generally, this model suggests that any perturbations that disrupt cell cycling and ultimately prevent the transit of cells through late G2 will also impair TMEJ. Consistent with this, similar to DNA2 inhibition, ATR inhibition resulted in a slightly slower transit from S-phase entry to G2 phase in both RPE1 and U2OS cells (Supplemental Figure S6E). Akin to DNA2i, ATRi treatment significantly reduced TMEJ utilization (Figure 5D, Supplemental Figure S7C). While it is tempting to imagine a hands-on role for ATR kinase in TMEJ (and indeed we have not ruled out such a role), a more parsimonious explanation is that any source of replication stress is liable to decrease the apparent usage of TMEJ.

## Discussion

In this study we report that normal DNA2 function is required for DSB repair by TMEJ. However, we find that the documented role for DNA2 in DSB resection is likely not the primary mechanism through which the nuclease exerts its observed effect on TMEJ. Rather, acute depletion of DNA2 (or inhibition of its catalytic activity) causes replication stress that prevents progression of cells to the G2/M transition when TMEJ is licensed. Through our efforts to understand this effect, we uncover new evidence that the optimal window for TMEJ is during the period of mitotic commitment in late G2 rather than in mitosis. We expand on prior studies implicating PLK1 in licensing of TMEJ by placing its activation primarily downstream of Aurora Kinase A (but not CDK1). Taken together, our findings suggest that cells pivot to TMEJ just prior to entering mitosis in an attempt to resolve unrepaired DSBs. During mitosis, a subset of persistent DSBs is likely subjected to more mutagenic outcomes driven by TMEJ leading to templated insertions.

Our initial motivation to initiate this study was to interrogate the role that DSB end resection plays in TMEJ. While DNA2 has well-documented, direct roles in DSB resection both *in vivo* and *in vitro* [18, 28, 94], we observed counterintuitively that its depletion in U2OS cells slightly elevated DSB resection (Figure 2D). However, as DNA2 depletion also increases the fraction of cells in S/G2 (Figure 2F), we suspect that this underlies the overriding stimulatory effect on the resection activity at the population level. Further study will be needed to disentangle cell cycle effects to directly determine how much resection (and which resection pathway) mechanistically supports TMEJ. As we now understand that cells must arrive at the G2/M transition in order to license TMEJ, future efforts leveraging small molecule inhibitors of MRE11, DNA2 [50], and EXO1 [95] to acutely inhibit nuclease function thereby allowing a much shorter time between DSB induction and sample collection may be necessary to avoid confounding cell cycle effects.

In dissecting the mechanism by which the slowed S phase induced by loss of DNA2 leads to diminished TMEJ, we provide further support for recent publications that describe PLK1-dependent phosphorylation of Pol θ as a requirement for mitotic TMEJ [57, 59, 62]. We note, however, that in focusing on mitotic repair these studies often pre-arrest cells in mitosis at the time of DNA damage and hold them there, thereby missing upstream events. We do indeed observe that TMEJ depends on PLK1, as evidenced by the 77% reduction in TMEJ:NHEJ ratio in asynchronous cells recovered into PLK1i for 24 hours (Figure 4B). Using a variety of additional conditions, we were able to more precisely examine the cell cycle timing of TMEJ’s activation. Delivering the Cas9 RNP immediately after release from PLK1i treatment leaves a sufficient window during which TMEJ can occur, leading to a far milder reduction in TMEJ:NHEJ ratio (28% reduced compared to untreated cells). Conversely, CDK1i addition at the time of RNP transfection does not significantly alter TMEJ usage. However, release from CDK1i treatment during RNP transfection more severely impairs TMEJ:NHEJ ratio (51% reduced compared to untreated cells) implying that cells synchronized at the G2/M transition with CDK1i have already missed the window when TMEJ is highly active. Mimicking the effects of CDK1i treatment, the addition of nocodazole does not impair TMEJ, but release out of a nocodazole block does impair TMEJ. These experiments collectively support a model where the majority of TMEJ repair events occur downstream of PLK1 activation but upstream of CDK1 activation and mitotic spindle assembly.

Even though CDK1i or nocodazole treatment does not alter bulk TMEJ usage, we were nevertheless intrigued by their inhibitory effects on templated insertions. We infer that both CDK1i and nocodazole treatments leave cells fully capable of engaging Polθ’s ability to anneal microhomologies and prime new synthesis from them, but unable to form secondary structures or perform the unwinding of new duplex DNA necessary for templated insertion formation. It has been demonstrated that Polθ requires both its polymerase domain and helicase-like domain to perform templated insertions, whereas simple deletion events mediated by pre-existing microhomologies require only the polymerase domain [96, 97]. We suspect this explains why Polθ in the presence of the polymerase-domain inhibitor ART558, which leaves its ATPase activity intact, can still drive templated insertions in RPE1 cells (Figure 1D). Similarly, *in vitro*, the polymerase domain is sufficient to extend simple microhomologies, but the helicase domain is needed for longer ssDNA substrates [10]. Biochemically, the Polθ helicase-domain facilitates ATP-independent ssDNA annealing [98–100], yet also possesses limited ATP-dependent DNA unwinding activity [101] and RPA displacement activity [99, 102]. These ATP-dependent functions would specifically support the mechanics of complex templated insertion events by first evicting RPA from the 3’ ssDNA tail to enable formation of secondary structures (e.g. hairpins) and then by unwinding the nascent duplex after Polθ terminates synthesis to permit a second round of annealing. Based on these insights, we propose that the range of Polθ-driven DSB repair outcomes that occur upon CDK1i and nocodazole treatment in mitosis may closely resemble that of ATPase-dead Polθ. While additional studies will be necessary to test this model, we hypothesize that during mitotic commitment prior to the G2/M transition the Polθ polymerase function is licensed by PLK1 while the activation of its ATP-dependent helicase activity occurs later in mitosis in a CDK1-dependent manner. While all reported PLK1 phosphorylation sites on Polθ reside in its central linker region [57], we note that the central domain is important for TMEJ repair *in vivo* and may modulate interactions between the polymerase and helicase domains [103]. Another mechanism may involve phosphorylation of the protein RHINO by CDK1 (and PLK1) driving its interaction with Polθ to DSBs that persist into mitosis [59]. We are also intrigued by reports of mitosis-specific phosphorylation of RPA2 at Ser-23 and Ser-29 by CyclinB-CDK1 that coincides with its exclusion from chromosomes [104], potentially enabling access of Polθ to longer ssDNA tails. Independent and temporal control of Polθ’s polymerase and ATPase activity may be advantageous to first drive the rapid resolution of simple, compatible DSB ends prior to mitosis; thereafter the processing of more complex DSBs would ensue in mitosis (i.e. repair of long RPA-coated ssDNA tracts without a nearby sister chromatid). Indeed, there is recent evidence that Polθ may also be recruited to collapsed replication forks during S and G2 [105], although its activation could be prompted only when PLK1 activity rises during mitotic commitment.

While our study largely supports recent reports that TMEJ is inhibited during interphase and operates during mitosis [56, 57, 59, 62], we can now add nuance based on the observation that the majority of TMEJ repair in our DSB repair assay depends on PLK1 but not CDK1. Taken together, we suggest that TMEJ licensing begins in late G2 when PLK1 activity rises, starting anywhere from ∼ 1 to 5 hours prior to nuclear envelope breakdown [61, 90]. We note that this PLK1-dependent and CDK1-independent repair comprises mostly simple microhomology-mediated deletions; more complex templated insertions are in fact sensitive to CDK1i or nocodazole treatment. We therefore reason that templated insertions likely occur after prometaphase, and that this repair requires activation of Polθ’s ATP-dependent helicase domain for either dsDNA unwinding or RPA displacement. For the specific Cas9-induced DSB used here these templated insertions are a minority of products (∼2.5% of total indels), but may nonetheless be of consequence. For studies of mitotic TMEJ that used genome-wide and nonspecific DNA damage (replication stress, ionizing radiation, topoisomerase poisons)[56, 57, 59], it is possible those more complex DNA lesions required full use of Polθ’s polymerase and helicase functions, thereby pushing the bulk of observable TMEJ repair further into mitosis.

There remains much to investigate about Polθ’s regulation. In particular, we do not know whether CDK1 has a direct role in facilitating templated insertions, or if its activity simply permits the requisite passage through mitosis. Phosphorylation by CDK1 often primes subsequent phosphorylation by PLK1 [87, 106] but whether this occurs with Polθ as a direct target remains unclear. Also, while roles for CDK1 in TMEJ are demonstrated here and elsewhere [59], it is unclear whether this occurs in complex with Cyclin A or B. We are also curious how drugs that disrupt the cell cycle may modulate TMEJ usage in tumors. There are numerous clinical trials in oncology for inhibitors of PLK1 [107], CDK1 [108], AURKA [109], and ATR [110]. We have demonstrated here that these inhibitors can impair TMEJ repair to varying degrees. It will be important to understand whether these drug treatments do indeed impair TMEJ in representative cancer models, and, if so, whether a TMEJ defect contributes to observed anti-tumor activity thereby offering a new therapeutic vulnerability.

## Materials and Methods

### Cell culture and siRNA treatments

RPE1-hTERT cells were cultured in DMEM/F12 (Gibco) supplemented with 10% fetal bovine serum (Gibco). All U2OS lines were cultured in DMEM (Gibco) supplemented with 10% fetal bovine serum (Gibco). All cells were maintained at 37°C in 5% CO_2._ RPE1 p53-knockout cells (RPE1 FRT/TR *ΔTP53*) were described in Tsukada *et al.* 2024 [111].

All siRNA transfections were performed according to manufacturer recommendations for the Dharmafect reagent. At the six-well plate scale, final concentrations were 25 nM siRNA and 1:400 Dharmafect 1 reagent (T-2001-03, Horizon Discovery). The following siRNAs from the Dharmacon ON-TARGETplus product line were used: Non-targeting siRNA #1 (D-001810-01-05), BLM (J-007287-06-0005), EXO1 (L-013120-00-0005), DNA2 (L-026431-01-0005), TOP3α (L-005279-00-0005), RMI1 (L-014527-01-0005). Cells were passaged as needed at 24 hours to maintain low confluency and adequate space for continued cell cycling. All experimental perturbations were performed 48 hours after siRNA transfection unless otherwise specified.

### CRISPR-Cas9 transfections

All site-specific DSBs were induced by electroporating a Cas9 ribonucleoprotein (RNP) into RPE1 or U2OS cells against the target sequence 5’ TACGCCACTGAACAGAACAA 3’. An Alt-R CRISPR-Cas9 crRNA XT (Hs.Cas9.ADGRL2.1.AG, IDT) and tracrRNA were hybridized, then complexed with S.p. Cas9 Nuclease V3 according to IDT recommendations.

Cells were electroporated using the Lonza 4D Nucleofector device according to manufacturer instructions for the P3 Primary Cell kit (RPE1) or SE Cell Line kit (U2OS). Final concentrations were 1 μM RNP and 1 μM Alt-R Cas9 Electroporation Enhancer (IDT) in each 100 μl electroporation reaction.

Electroporations were immediately recovered into fresh media and seeded to ∼ 50% confluence into either normal media or drug-treatment media. Cells were harvested 24 hours later for genomic DNA extraction unless otherwise specified.

### Resection assay

U2OS cells pre-treated with siRNAs and transfected with Cas9 RNP as above. Cells were harvested at an 8 hr timepoint, however, for genomic DNA extraction. The protocol for quantifying resected intermediates was adapted from Zhou, Y., *et al* 2014 [67]. This protocol exploits the fact that ssDNA (resected DNA) will be protected from cleavage by the ApoI restriction enzyme, but dsDNA (non-resected DNA) will not be.

Equal quantities of genomic DNA were subjected to an ApoI digest and a HincII digest. ApoI recognition sites are common; there is ApoI site 1040 bp away from the Cas9 cut site. HincII is used a mock digest as there are not any HincII sites within the regions being probed. Restriction digests were as follows: 20 μl total volume, ∼ 200 ng DNA, 1X NEB rCutSmart buffer, 10 U ApoI-HF (R3566S, NEB) or 10 U HincII (R0103S, NEB). The reactions were incubated at 37°C for 4 hours and heat-inactivated at 80°C for 20 minutes. Digests were then diluted to 50 μl with nuclease-free water. The diluted digests were then used as templates in qPCR reactions. Each 20 μl qPCR reaction consisted of: 2 μl template, 0.5 μM forward primer, 0.5 μM reverse primer, 1X qPCR iTaq Universal SYBR Green Supermix (1725121, Bio-Rad). Reactions were run in a StepOne Plus Real Time PCR System (Applied Biosystems) using software version 2.3. Cycling conditions were: initial denaturation at 95°C for 30 s, then 40X cycles of 95°C for 3 s, 60°C for 30 s.

Primers sets were designed to amplify across ApoI cut sites in the vicinity of the Cas9-induced DSB, or across an ApoI site in the ß-actin gene. The ß-actin ApoI site is not near the Cas9-induced DSB, and therefore remains double-stranded and should be fully cleaved by the ApoI digest. This acts as a control for the digestion efficiency. Primer sequences are included in Table S2.

Calculation are as follows: ΔCt = Ct_ApoI-digest_ – Ct_HincII-digest_. ssDNA% = 1/(2^(△Ct-1) + 0.5)*100.

### LMNA-Clover HDR reporter assay

U2OS cells were first siRNA-treated as described above for 72 hours. Cells were then collected and electroporated with an sgRNA and Cas9 expression plasmid (pX330-LMNA-gRNA1, Addgene #122507) and an mClover donor plasmid (pCR2.1 Clover-LMNA Donor, Addgene #122508) [76]. Each 100 μl electroporation reaction contained 1.5 μg pX330-LMNA-gRNA1 and 1 μg pCR2.1 Clover-LMNA Donor.

Cells were electroporated using the Lonza 4D Nucleofector device according to manufacturer instructions for the SE Cell Line kit (V4XC-1024, Lonza). Cells were recovered immediately into fresh media and re-seeded. After 72 hours of incubation, cells were harvested, resuspended in PBS, and analyzed for mClover expression by flow cytometry on a Cytoflex LX instrument (Beckman Coulter). Fluorescence signal in the FITC channel was recorded for roughly 30,000 live, single cells. Gating was performed on the FITC-A histogram of untransfected cells from each siRNA-treatment condition such that 0.25% of cells were assigned FITC-positive. This gate was then applied to the transfected samples and the 0.25% ‘false positive’ rate was subtracted to yield % mClover-positive.

### NGS library preps and indel analysis

Genomic DNA was used a template for a PCR reaction that spanned Cas9 cut site. Forward and reverse primers contained 5’ extensions with unique 7-bp barcode sequences (Table S3). Each genomic DNA samples within a library was amplified with a unique primer combination for identification. Each 50 μl reaction contained ∼ 200 ng gDNA, 0.5 μM forward primer, 0.5 μM reverse primer, and Q5 Hot Start High-Fidelity Master Mix at 1X (M0494S, NEB). Cycling conditions were: initial denaturation at 98°C for 30 s, then (98°C for 10 s, 66°C for 15 s, 72°C for 30 s) 35X cycles, 72°C for 2 min.

All PCR reactions within a given library were then pooled equally and purified together with the Monarch PCR and DNA Cleanup Kit (T1030L, NEB). The purified PCR products were then library prepped with the NEBNext Ultra™ II DNA Library Prep Kit (E7103S, NEB) and indexed with NEBNext Multiplex Oligos for Illumina (E6440S, NEB) according to the manufacturer’s protocol. Briefly, the amplicons in a library were end-ligated with adaptor sequences, and then PCR-enriched to install i7 and i5 indices. All purification steps were performed with size-selection beads. Libraries were sequenced at the Yale Center for Genomic Analysis on a NovaSeq platform with 2×150 read length.

Combinatorial demultiplexing the paired-end reads is performed by Cutadapt (version 4.2) using an error rate of 1. Repair products were classified into types (insertions, deletions, SNV, templated insertions, etc) using the computational tool Sequence Interrogation and Quantification (SIQ, version 1.3) [70]. Mutation data were visualized in SIQPlotteR. Deletions with a microhomology ≥ 2 were scored as TMEJ, whereas +1 bp insertion or deletions with microhomology ≤ 1 were scored as NHEJ. Raw data will be available under BioProject submission SUB14755135.

### Cell cycle analysis flow cytometry

Unless otherwise specified, all cell cycle analysis was performed on cells 48 hours after siRNA transfection or 24 hours after drug treatment. Cells were harvested from a ∼50% confluent 10cm tissue culture dish, washed once in PBS, and resuspended in 100 μl PBS. Cells were then dispensed drop-wise into 3 mL of pre-chilled 70% ethanol in a 15 mL conical tube while vortexing. Tubes were kept on ice for 15 minutes. Cells were then pelleted, washed once in PBS, and resuspended in 500 μl propidium iodide (PI)/RNaseA staining buffer (550825, BD).

DNA content was measured by flow cytometry. PI signal was recorded in the PE-TexasRed channel of a Cytoflex LX instrument (Beckman Coulter). Cell cycle distributions were determined using the Watson (Pragmatic) model in FlowJo v10.10.0.

### Cell viability assays

Cells were seeded in a 96-well plate at 1000 cells/well or 24 -well plates at 10,000 cells/well. If a pre-treatment was required, cells were first siRNA-treated for 48 hours prior in 6-well or 10cm plates before seeding. Drug treatments were left to incubate for 24 hours before being washed out. After roughly 5 days (before any wells became fully confluent) media was exchanged with a 1:1 mix of serum-free media and 5 mg/mL MTT (M6494, Invitrogen) in PBS. Cells with left to incubate at 37°C for 4 hours, after which all MTT solution was removed and replaced with DMSO for 30 minutes. Absorbance at 560 nm was recorded for each well on a Biotek Synergy 2 plate reader. After background subtraction, absorbance values were normalized to the average untreated absorbance to derive relative viability. For one assay, crystal violet staining was instead used. Briefly, cells were stained with a crystal violet solution (0.25% crystal violet, 3.5% formaldehyde, 72% methanol), then rinsed and dried. The crystal violet was dissolved in a 10% acetic acid solution and absorbance at 590 nm was measured on a plate reader.

### qRT-PCR

RNA was extracted after 48 hours of siRNA knockdown using the Monarch Total RNA Miniprep Kit (T2010S, NEB). RNA was reverse transcribed using the High-Capacity cDNA Reverse Transcription Kit (4368814, Applied Biosystems) using the included random primers in a 20 μl. The cDNA was diluted to 50 μl with nuclease-free water and used as a template for qPCR. Each 20 μl qPCR reaction consisted of: 2 μl template, 0.5 μM forward primer, 0.5 μM reverse primer, 1X qPCR iTaq Universal SYBR Green Supermix (1725121, Bio-Rad). Reactions were run in a StepOne Plus Real Time PCR System (Applied Biosystems). Cycling conditions were: initial denaturation at 95°C for 30 s, then 40X cycles of 95°C for 3 s, 60°C for 30 s. Relative mRNA expression was determined using the ΔΔCt method with an *HPRT1* reference.

## Acknowledgements

We thank all members of the Jensen and LusKing labs for invaluable insights and discussions. We thank Diane King and Sunnycrest Bioinformatics for developing the computational analysis of the sequencing data. RPE1 FRT/TR *ΔTP53* cells were generously provided by Dr. Andrew Blackford. This research was supported by grants from the NIH (R01 CA215990, R01 CA270788), a Research Scholar Grant from the American Cancer Society, and the Women’s Health Research at Yale (to R.B.J.), the Gray Foundation (to R.B.J. and M.C.K.), and Pilot Funding from the Yale Cancer Center (to M.C.K., P30 CA016359). C.P.M. was also supported by T32GM007223. Research reported in this publication was supported by the National Institute of General Medical Sciences of the National Institutes of Health under Award Number 1S10OD030363-01A1. We thank Yale Flow Cytometry for their assistance. The Core is supported in part by an NCI Cancer Center Support Grant P30 CA016359.

