## Supplemental Figures S1-S9 for "Activation of Pol theta-mediated end joining begins during mitotic commitment"

### Supplement S1

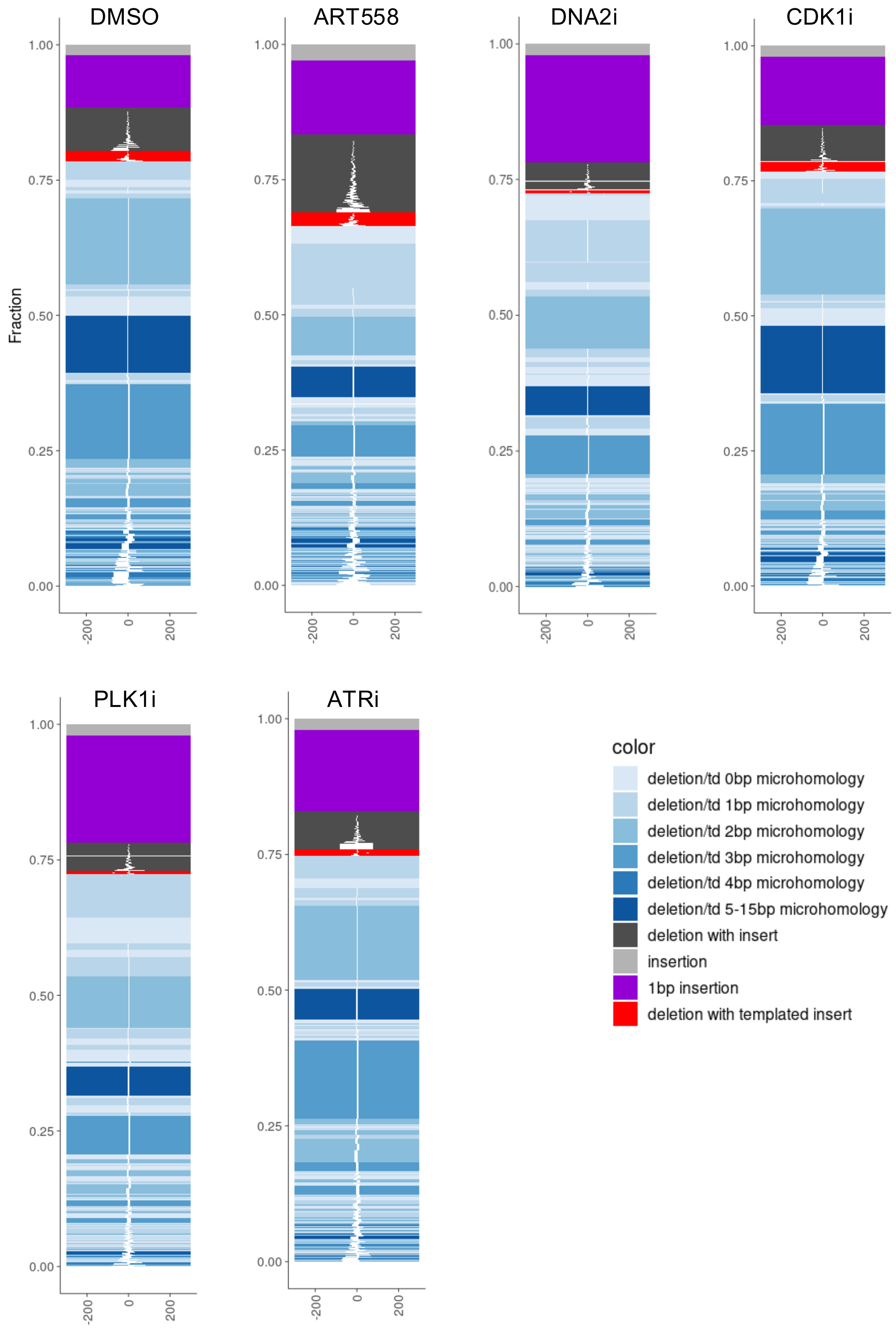

**Supplement S1.** Representative repair spectra of RPE1 cells incubated with the indicated drug treatments for 24 hours after DSB induction by Cas9. The plots selected have the median TMEJ:NHEJ ratio for the given drug treatment.

### Supplement S2

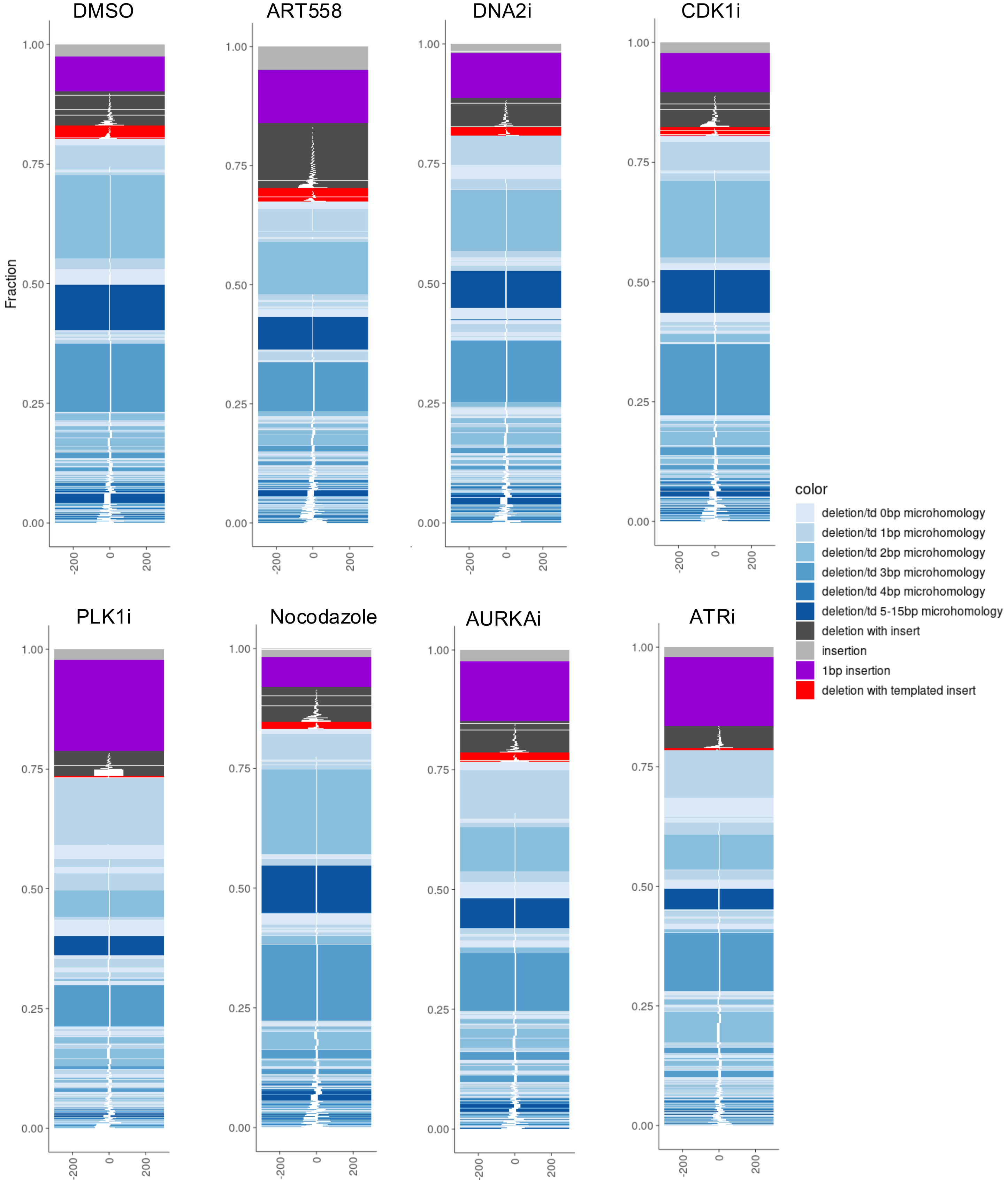

**Supplement S2.** Representative repair spectra of U2OS cells incubated with the indicated drug treatments for 24 hours after DSB induction by Cas9. The plots selected have the median TMEJ:NHEJ ratio for the given drug treatment.

### Supplement S3

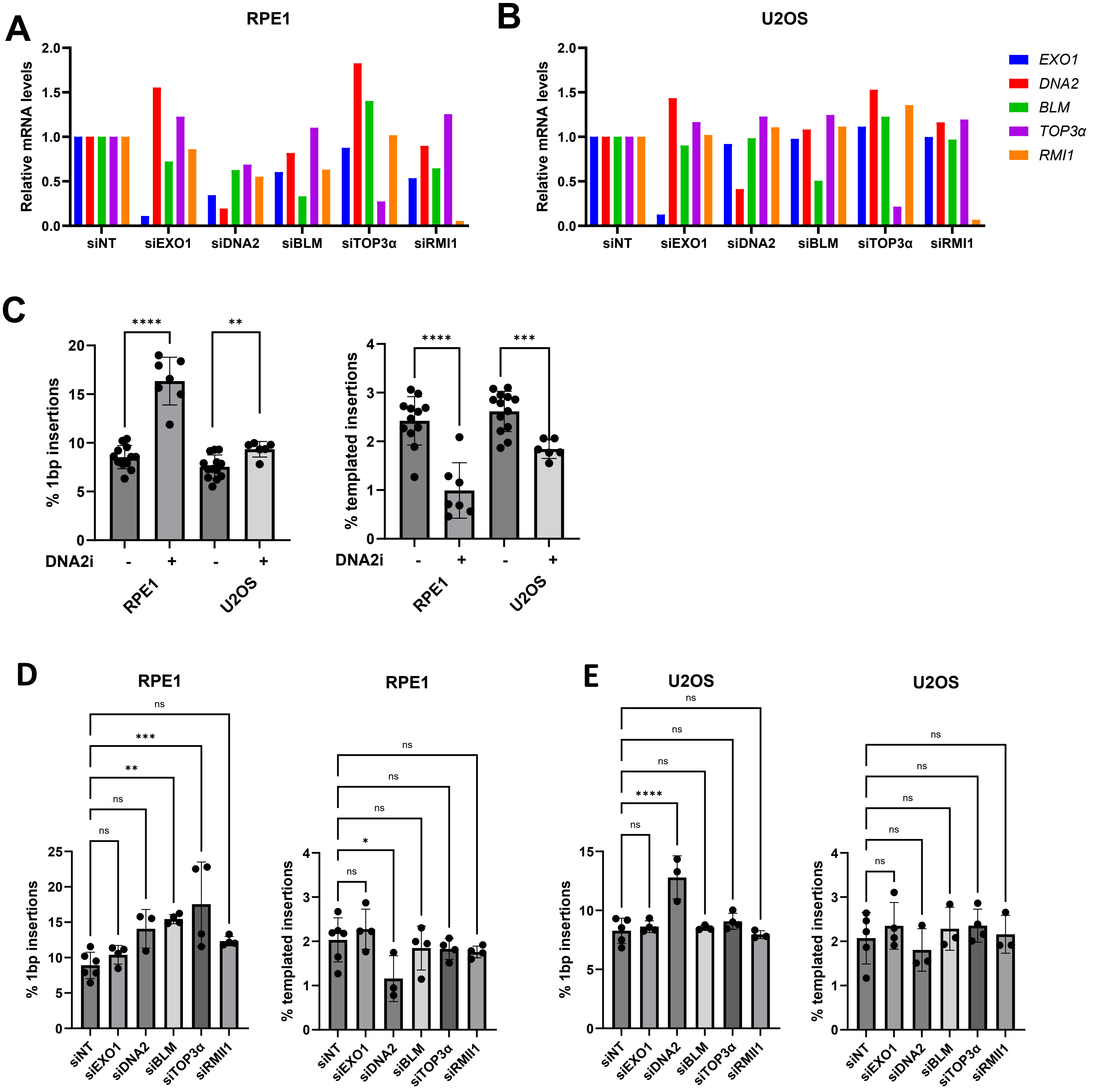


**Supplement S3. (A-B)** Relative expression levels of target genes after 48 hours of siRNA knockdown in **(A)** RPE1 cells and **(B)** in U2OS cells. **(C)** Percentage of 1bp insertions and templated insertions (as a % of total indels) for RPE1 and U2OS cells treated with the DNA2 inhibitor C5 (60 µM) for 24 hours post Cas9 RNP transfection. Data are mean ± SD of at least 6 independent biological replicates. Significance for each pair was determined by unpaired, two-tailed t-test. **(D-E)** Percentage of 1bp insertions and templated insertions (as a % of total indels) for the indicated siRNA treatments in **(D)** RPE1 and **(E)** U2OS cells. Data are mean ± SD of at least three independent biological replicates. Significance was determined by one-way ANOVA with Šídák's multiple comparison test.

### Supplement S4

A

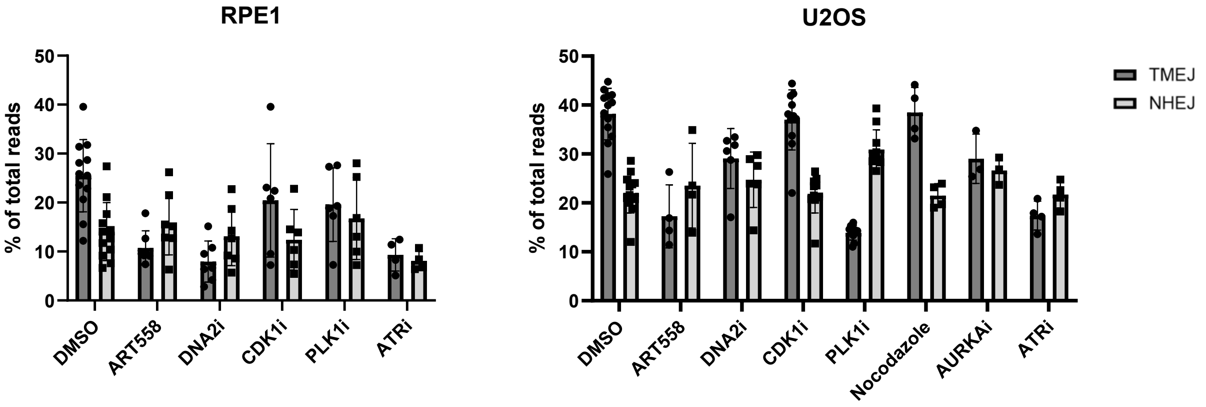


B

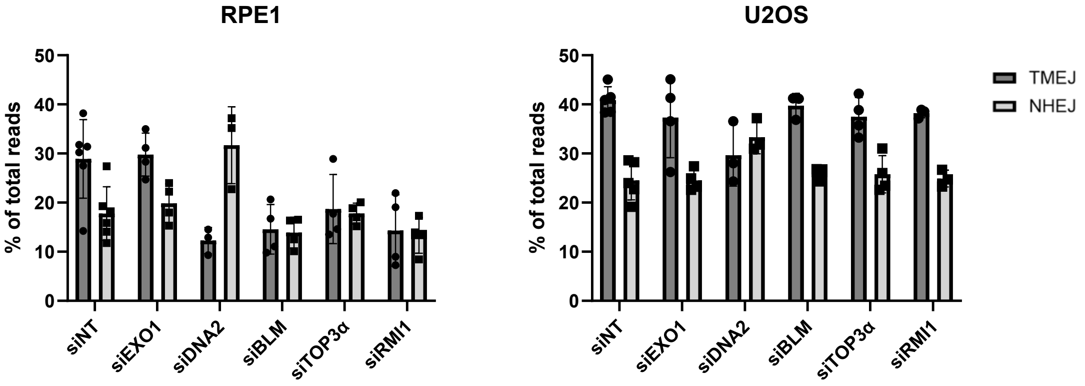


**Supplement S4.** Summary of RPE1 and U2OS cells transfected with the Cas9 RNP and **(A)** recovered into the indicated drug treatment or **(B)** pre-treated with the indicated siRNAs depicting the raw percentage of TMEJ events vs NHEJ events. **(A)** 10 µM ART558, DNA2i = 60 µM C5, CDK1i = 5 µM RO-3306, PLK1i = 0.1 µM BI-2536, 0.2 µM Nocodazole, AURKAi = 1 µM Alisertib, ATRi = 5 µM VE-822 in RPE1, 0.5 µM VE-822 in U2OS.

### Supplement S5

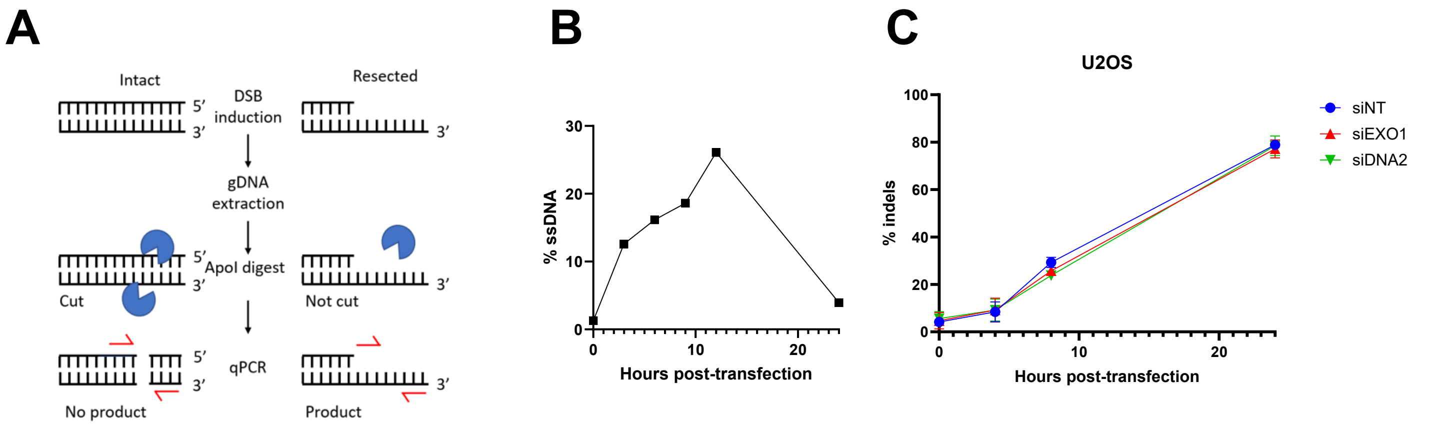


**Supplement S5. (A)** Schematic of the qPCR-based resection assay. Genomic DNA harvested at various times after Cas9 RNP transfection. Purified gDNA is digested *in vitro* with ApoI to degrade dsDNA. Uncut ssDNA is quantified by qPCR. **(B)** Representative time course of resection 1040 bp distal to the Cas9-induced DSB. **(C)** Total percent indels at various timepoints in U2OS cells siRNA-depleted of the indicated genes. Data are mean ± SD, n=2 independent biological replicates.

### Supplement S6

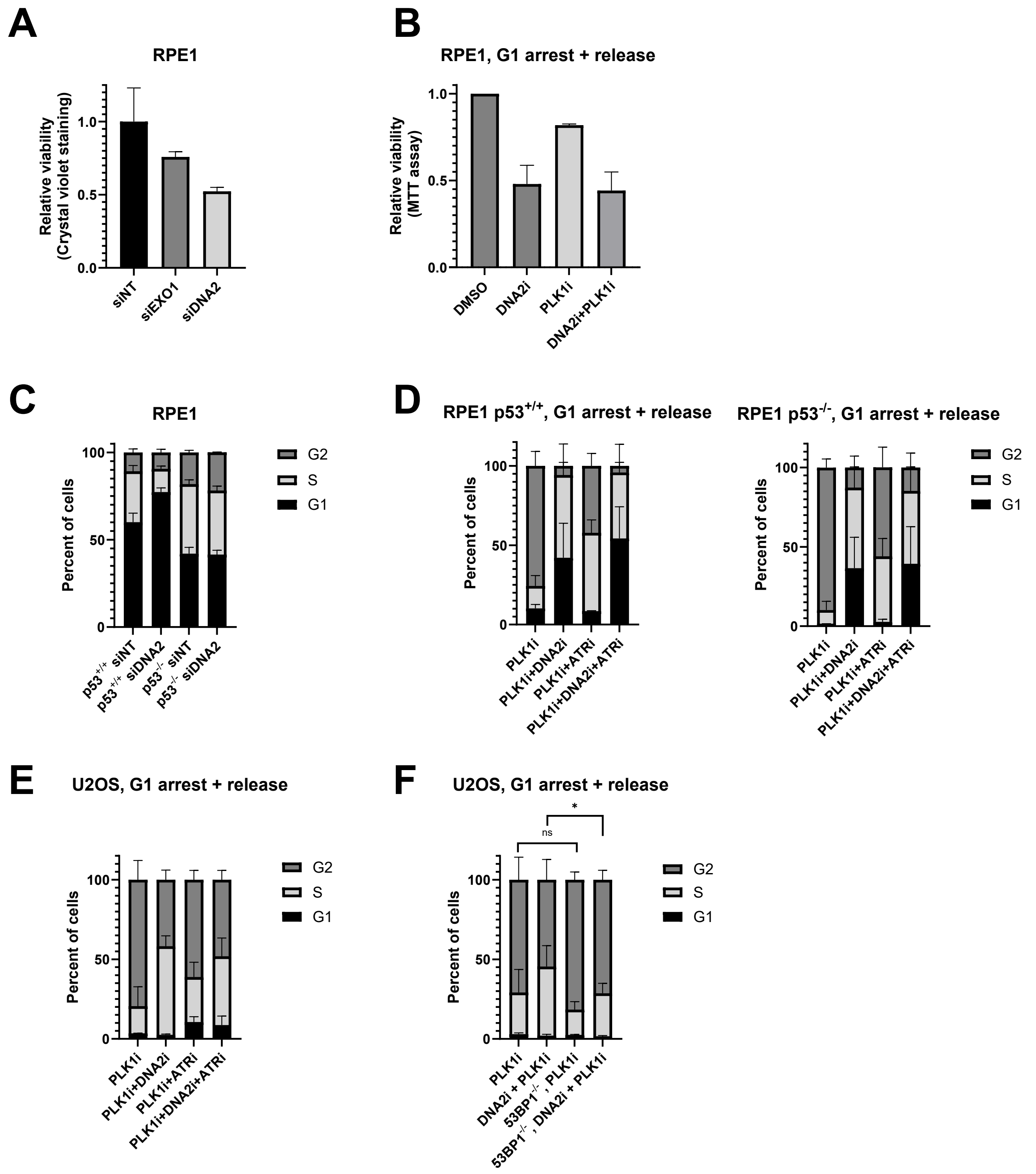


**Supplement S6. (A)** Relative viability of RPE1 cells treated with the indicated siRNAs and left to outgrow for 4 days. Viability was assessed by crystal violet staining. Data are mean ± SD, n=2 technical replicates. **(B)** Relative viability of RPE1 cells which were serum-starved for 24 hours, then released into complete media with the indicated drug combinations for 24 hours. Cells were left to outgrow for 5 days in untreated media before viability assessment with an MTT assay. Data are mean ± SD, n =2 independent biological replicates. **(C)** Cell cycle analysis of p53-proficient and deficient RPE1 cells treated for 48 hr with the indicated siRNAs. Data are mean ± SD, n=3 independent biological replicates. **(D)** Cell cycle analysis of RPE1 $p53^{+/+}$ and $p53^{-/-}$ cells serum-starved for 24 hours and released into complete media containing the indicated drug combinations for 24 hours. Data are mean ± SD, n =2 independent biological replicates. **(E)** Cell cycle analysis of U2OS cells serum-starved for 24 hours and released into complete media containing the indicated drug combinations. Data are mean ± SD, n=2 independent biological replicates. **(F)** U2OS with and without 53BP1 serum-starved and released into complete media with the indicated drug combinations. Data are mean ± SD, n = at least 5 independent biological replicates. Significance was determined by one-way ANOVA with Šídák's multiple comparison test **(B-F)** DNA2i = 60 µM C5, PLK1i = 0.1 µM BI-2536, ATRi = 5 µM VE-822 in RPE1, 0.5 µM VE-822 in U2OS. * $p \leq 0.05$, ns = not significant.

### Supplement S7

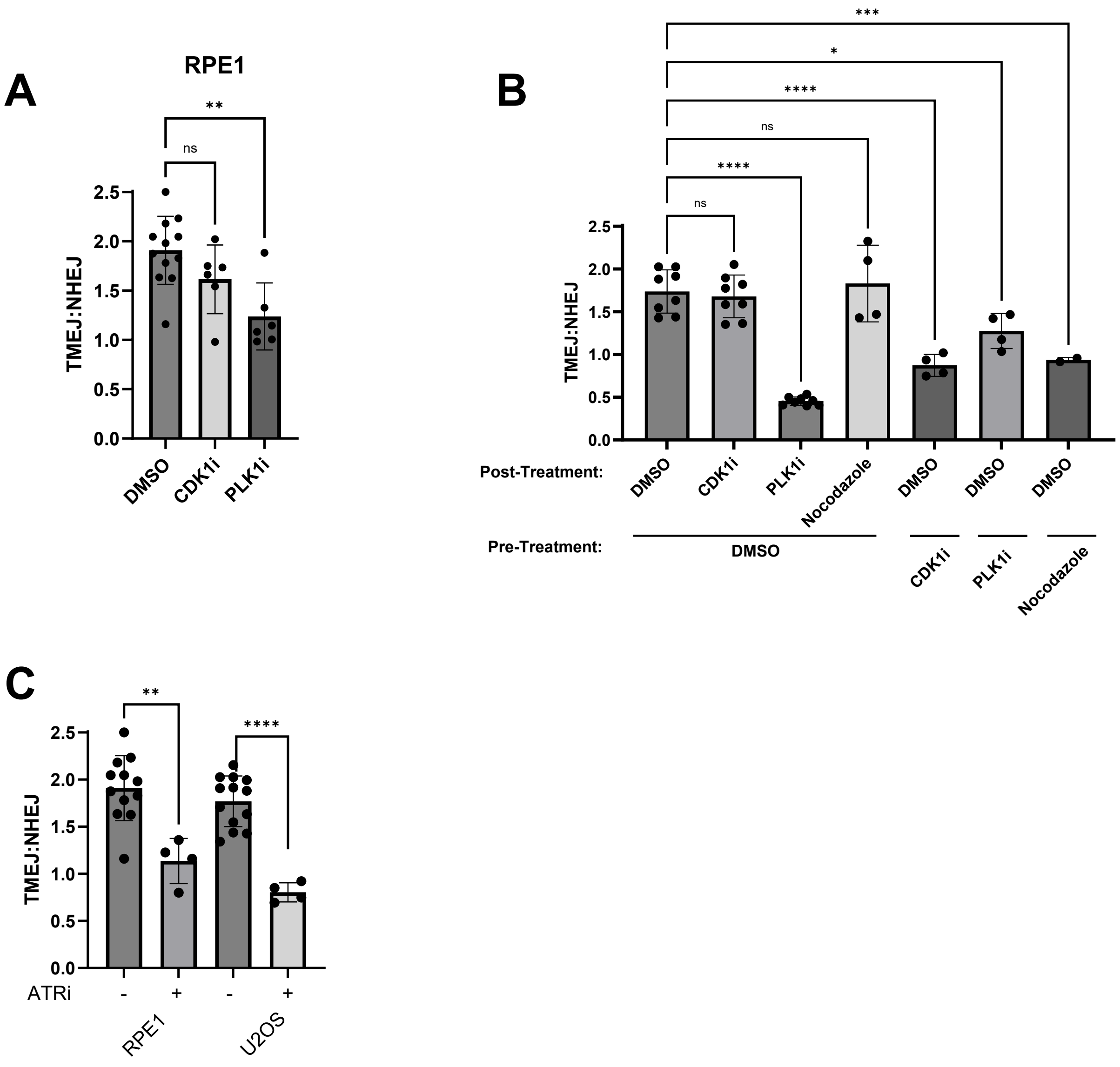


**Supplement S7. (A)** TMEJ vs NHEJ repair events of RPE1 cells cultured asynchronously and recovered into the indicated drug treatments after Cas9 RNP transfection. Data are mean ± SD of at least 6 independent biological replicates. Significance was determined by one-way ANOVA with Šídák's multiple comparison test. **(B)** TMEJ vs NHEJ repair events of U2OS cells either cultured asynchronously for 24 hours or arrested at the G2/M transition with CDK1i, PLK1i, or Nocodazole treatment prior to Cas9 RNP transfection (Pre-Treatment) . After transfection, cells were recovered into DMSO, CDK1i, or PLK1i treatments (Post-Treatment). Data are mean ± SD of at least 2 independent biological replicates. Significance was determined by one-way ANOVA with Šídák's multiple comparison test. **(C)** TMEJ vs NHEJ repair events of a Cas9-induced DSB in RPE1 and U2OS cells recovered into ATRi for 24 hours. Significance for each pair was determined by unpaired, two-tailed t-test. **(A-C)** DNA2i = 60 μM C5, CDK1i = 5 μM RO-3306, PLK1i = 0.1 μM BI-2536, ATRi = 5 μM VE-822 in RPE1, 0.5 μM VE-822 in U2OS. * $p \leq 0.05$, ** $p \leq 0.01$, *** $p \leq 0.001$, **** $p \leq 0.0001$, ns = not significant.

### Supplement S8

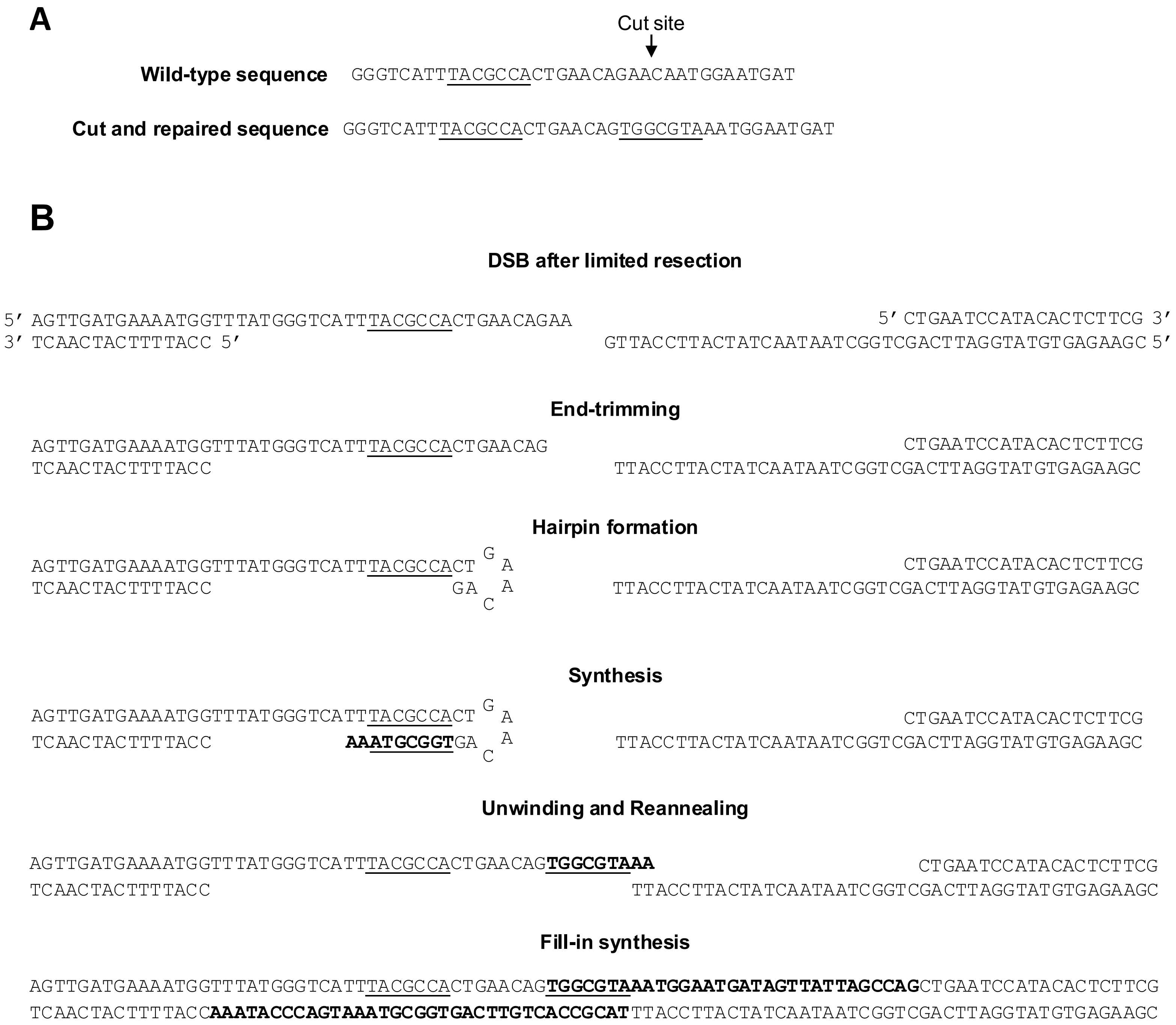


**Supplement S8. (A)** The uncut, wild-type sequence used for the DSB repair assay and a templated insertion repair event which is highly represented in our sequencing datasets. Insertions arising from TMEJ repair are often homologous to another sequence closely flanking the cut site. In this example, the underlined sequence present in the uncut sequence is inserted in the reverse complement orientation (inverted repeat) during TMEJ repair. **(B)** A putative mechanism for this templated insertion demonstrates the various steps required. In this example, only short-range resection of the DSB is drawn, though much longer resection tracts and long ssDNA tails coated by RPA are probable *in vivo*. New synthesis is depicted in bold.

#### Supplement S9

**A**

##### RPE1

| | DMSO | ART558 | DNA2i | CDK1i | PLK1i | ATRi |
|---|---|---|---|---|---|---|
| **TMEJ:NHEJ** | **1** | **0.382** | **0.319** | **0.846** | **0.649** | **0.596** |
| del MH ≥ 2bp | 1 | 0.529 | 0.569 | 0.957 | 0.858 | 0.784 |
| del MH ≤ 1bp | 1 | 1.360 | 1.761 | 1.049 | 1.237 | 1.160 |
| 1bp insertion | 1 | 1.432 | 1.916 | 1.334 | 1.560 | 1.637 |
| TINS | 1 | 1.306 | 0.409 | 0.536 | 0.599 | 0.774 |
| DELINS | 1 | 2.005 | 0.635 | 0.806 | 0.856 | 0.888 |

**B**

##### U2OS, from Figure 4B

| Pre-Treatment | DMSO | | | CDK1i | | | PLK1i | | |
|---|---|---|---|---|---|---|---|---|---|
| Post-Treatment | DMSO | CDK1i | PLK1i | DMSO | CDK1i | PLK1i | DMSO | CDK1i | PLK1i |
| **TMEJ:NHEJ** | 1 | 0.972 | 0.234 | 0.492 | 0.487 | 0.178 | 0.719 | 0.577 | 0.271 |
| del MH ≥ 2bp | 1 | 1.008 | 0.480 | 0.739 | 0.737 | 0.386 | 0.865 | 0.788 | 0.524 |
| del MH ≤ 1bp | 1 | 1.016 | 1.871 | 1.453 | 1.456 | 2.013 | 1.202 | 1.266 | 1.788 |
| 1bp insertion | 1 | 1.088 | 2.593 | 1.661 | 1.731 | 2.627 | 1.220 | 1.757 | 2.393 |
| TINS | 1 | 0.668 | 0.126 | 0.561 | 0.419 | 0.101 | 0.805 | 0.543 | 0.164 |
| DELINS | 1 | 1.007 | 0.610 | 0.846 | 0.842 | 0.603 | 1.078 | 0.975 | 0.680 |

**Supplement S9. (A-B)** Summary heatmap of DSB repair events in **(A)** RPE1 cells incubated with the indicated drug treatments for 24 hours after Cas9 RNP transfection and **(B)** U2OS cells from the experiment depicted in Figure 4B. For simpler comparison, all quantities are normalized to the DMSO average for the given mutation type/metric. The normalized TMEJ:NHEJ ratio and the three events that comprise it are boxed; TINS (templated insertions) and DELINS (deletions with insertions) are not incorporated into this ratio but are nevertheless highly informative. **(A-B)** 10 µM ART558, DNA2i = 60 µM C5, CDK1i = 5 µM RO-3306, PLK1i = 0.1 µM BI-2536, ATRi = 5 µM VE-822 in RPE1.
